# A novel Fis1 inhibiting peptide reverses diabetic endothelial dysfunction in human resistance arteries

**DOI:** 10.1101/2020.11.16.385054

**Authors:** Kelsey A. Nolden, Mamatha Kakarla, John M. Egner, Jingli Wang, Megan C. Harwig, Venkata K. Puppala, Benjamin C. Hofeld, Leggy A. Arnold, David Z. Trykall, Francis C. Peterson, Michelle L. Roberts, David M. Jenson, R. Blake Hill, Michael E. Widlansky

## Abstract

Mitochondrial dysfunction is one of several factors that drive development of vascular endothelial dysfunction in type 2 diabetes (T2DM). In endothelial cells from T2DM patients, mitochondrial networks are highly fragmentated with increased expression of mitochondrial fission protein 1 (Fis1). However, whether manipulation of Fis1 expression and activity in endothelial vessels from T2DM patients alters endothelial function remains unknown. Here, molecular suppression of Fis1 reversed impaired endothelium-dependent vasodilation of vessels from T2DM patients, as well as healthy human vessels exposed to high (33 mM) or low (2.5 mM) glucose, while preserving NO bioavailability and improving endothelial cell layer integrity. Conversely, overexpression of Fis1 in healthy vessels impaired vasodilation and increased mitochondrial superoxide, suggesting a causative role. Application of a novel and specific Fis1 inhibitor, pep213, improved endothelium-dependent vasodilation of vessels from T2DM patients, as well as healthy vessels exposed to high glucose or Fis1 overexpression, by improving NO bioavailability and decreasing excess mitochondrial ROS generation. The specificity of pep213 was determined through multiple biophysical techniques and a 1.85 Å crystal structure of pep213 in complex with Fis1. These data support that excessive mitochondrial fragmentation drives endothelial vessel dysfunction and supports a potential novel therapeutic route for treating diabetic microvascular disease through pharmacological inhibition of Fis1.

## Introduction

Vascular endothelial dysfunction is a well-established precursor to the development of both microvascular and macrovascular disease in patients with type 2 diabetes mellitus (T2DM) (Brownrigg *et al*, 2016; Halcox *et al*, 2002; Mohammedi *et al*, 2017). Several factors drive endothelial dysfunction including oxidative stress from NADPH oxidase, increased flux through the polyol pathway, increased protein glycation, and aberrant activation of redox-sensitive protein kinase (Barrett *et al*, 2017). A common feature of these pathological drivers is mitochondrial dysfunction with elevated reactive oxygen species (ROS) that contributes to pathogenesis. In liver epithelial cells from rats fed high glucose, mitochondria become hyperpolarized, driving ROS formation that is dependent upon fragmentation of mitochondrial networks (Yu *et al*, 2006). In endothelial cells from individuals with T2DM, similar phenomena are observed with increased polarization of the mitochondrial inner membrane, hyper-fragmented networks (Shenouda *et al*, 2011), and increased ROS (Kizhakekuttu *et al*, 2012; Nishikawa *et al*, 2000). Elevated ROS formation correlates with increased apoptosis (Yu *et al*, 2008) and activates deleterious epigenetic changes and cell signaling pathways that are thought to drive inflammation and vascular dysfunction (El-Osta *et al*, 2008). These cell-based studies support that controlling mitochondrial ROS by attenuating either mitochondrial membrane potential or network fragmentation during hyperglycemia may be of potential benefit.

Towards this goal, mitochondrial-targeted antioxidants or pharmacological agents that partially depolarize the mitochondrial inner membrane can reverse impaired endothelium-dependent vasodilation in resistance arterioles from humans with T2DM (Kizhakekuttu *et al*, 2012; Wang *et al*, 2012). Unfortunately, phase 3 clinical trials of therapeutic antioxidant approaches to prevent and treat vascular diseases have failed to recapitulate the positive effects seen in animal, epidemiologic, and non-randomized studies (Yusuf *et al*, 2000; Willcox *et al*, 2008); and, current pharmacological agents that target the mitochondrial inner membrane have toxicity profiles that preclude their clinical use (Widlansky & Hill, 2018). Therefore, targeting proteins involved in hyperglycemia-induced mitochondrial fragmentation presents a promising alternative approach. Prior work in venous endothelial cells from T2DM patients found increased expression of mitochondrial fission proteins Fis1 and Drp1, and decreased expression of the mitochondrial fusion protein MFN2. Further, RNAi-based knockdown of Fis1 or Drp1 was found to block hyperglycemic-induced increases in mitochondrial superoxide production and impairments in endothelium-derived nitric oxide synthase (eNOS) activation in endothelial cells (Shenouda *et al*, 2011). For Drp1, these cell-based findings directly translated to resistance arterioles from T2DM patients in *ex vivo* experiments where siRNA of Drp1 improved poor vasodilation (Tanner *et al*, 2017). These experiments support the benefit of blocking mitochondrial fragmentation, but the occurrence of several Drp1 pathological variants that cause neurological disorders and neonatal lethality raise questions about the therapeutic window for Drp1 inhibition.

By contrast to Drp1, no pathological variants have been identified for Fis1 to date. Fis1 may also be a more attractive pharmacological target given its role in stress-induced, not homeostatic, fission (Kleele *et al*, 2021). Indeed, responses to pathological stimuli like hypoxemia, hyperglycemia, infection, and oxygen deprivation appear to be mediated by Fis1 (Kim *et al*, 2011; Kumari *et al*, 2012; Ciarlo *et al*, 2012; Haileselassie *et al*, 2019, 2020; Disatnik *et al*, 2013). The Fis1-Drp1 axis also appears to be involved in a variety of neurodegenerative diseases such as Huntington’s disease, amyotrophic lateral sclerosis, and Alzheimer’s disease, where inhibition of the Fis1-Drp1 interaction appears very promising (Guo *et al*, 2013; Joshi *et al*, 2018a, 2018b, 2019; Qi *et al*, 2013). Here, we establish that overexpression of Fis1 in small resistance vessels from healthy volunteers diminishes their ability to vasodilate normally, whereas molecular suppression of Fis1 rescues impaired vasodilation in vessels under both hyper- and hypoglycemic conditions, as well as vessels from patients with T2DM, suggesting a causative role. Consistent with this idea, a novel and specific inhibitor of Fis1, pep213, was also found to improve impairments in endothelium-dependent vasodilation. These data support a critical role for Fis1 in the development of diabetic endothelial dysfunction and suggest that pharmacological targeting of Fis1 may hold promise for treating vascular disease in T2DM.

## Results

### Suppressing Fis1 improves endothelium-dependent vasodilation and increases NO bioavailability in human vessels

Impaired microvascular endothelial function is a well-known predictor of future adverse cardiovascular events, even in the absence of concomitant epicardial coronary artery disease (Brownrigg *et al*, 2016; Mohammedi *et al*, 2017; Halcox *et al*, 2002). To study the role of Fis1 in endothelial vessel vasodilation, we used small resistance arterioles from gluteal fat pad biopsies of normal and T2DM volunteers as well as from discarded subcutaneous adipose tissue from operative procedures (**Figure 1A**). We first confirmed the poor vasodilatory response upon acetylcholine challenge from T2DM compared to healthy vessels (**Figure 1B**). As vessels from normal volunteers were more plentiful, we next asked if molecular suppression of Fis1 by a specific siRNA incubated with normal vessels exposed to high (**Figure 1C**; 33 mM) or low (**Figure 1D**; 2.5 mM) glucose, equivalent to 594 and 45.05 mg/dL respectively, had an impact on vasodilation. Fis1 siRNA treatment reversed impaired vasodilation compared to a control siRNA treatment under as determined by a two-way ANOVA with post hoc analysis using Dunnett’s multiple comparison test at the highest concentration of acetylcholine. These favorable effects were completely attenuated by concurrent treatment with L-NAME, a potent endothelial nitric oxide synthase (eNOS) inhibitor (Rees *et al*, 1990), confirming these effects are endothelial-dependent. Fis1 siRNA and Fis1 over-expression negligible effects on vascular smooth muscle reactivity as measured by papaverine exposure. (**Supplemental Figure 1A**). Increasing Fis1 expression nearly 11-fold (**Supplemental Figure 1B**) in normal vessels significantly decreased vasodilation (**Figure 1E**). Treatment of healthy vessels with Fis1 siRNA did not affect vasodilation (**Figure 1F**), consistent with the idea that Fis1 is pathologically upregulated upon exposure to high or low glucose. We next asked whether suppression of Fis1 in vessels from patients with T2DM improved vasodilation. Treatment with Fis1 siRNA, decreased protein levels by ∼50% (**Supplemental Figure 1C**) and improved endothelium-dependent vasodilation (**Figure 1G**), although with a 14% lower maximum vasodilatory capacity compared to healthy vessels with the same treatment (**Figure 1H**). Consistent with L-NAME inhibition of eNOS in these experiments, siRNA transfection of either Fis1 or Drp1 significantly increased NO availability in vessels from healthy individuals exposed to either low or high glucose (**Supplemental Figure 2; Supplemental Figure 3A-B**). Drp1 siRNA treatment also improved impaired vasodilation in healthy vessels exposed to high glucose (**Supplemental Figure 3C**), as well as trended towards improving vasodilation in T2DM vessels (**Supplemental Figure 3D**), further supporting the idea that inhibiting aberrant mitochondrial fission is beneficial. Suppressing Fis1 in endothelial cells also improved NO bioavailability and barrier function without perturbing cellular energetics (**Supplemental Figure 4**) or the expression of other proteins involved in mitochondrial dynamics, function, or autophagy (**Supplemental Figure 5**). These new findings indicate that Fis1 levels can govern endothelial-dependent vasodilation in an eNOS-dependent manner and support that pharmacological inhibition of Fis1 may improve vasodilation by increasing bioavailable nitric oxide.

**Figure 1.**
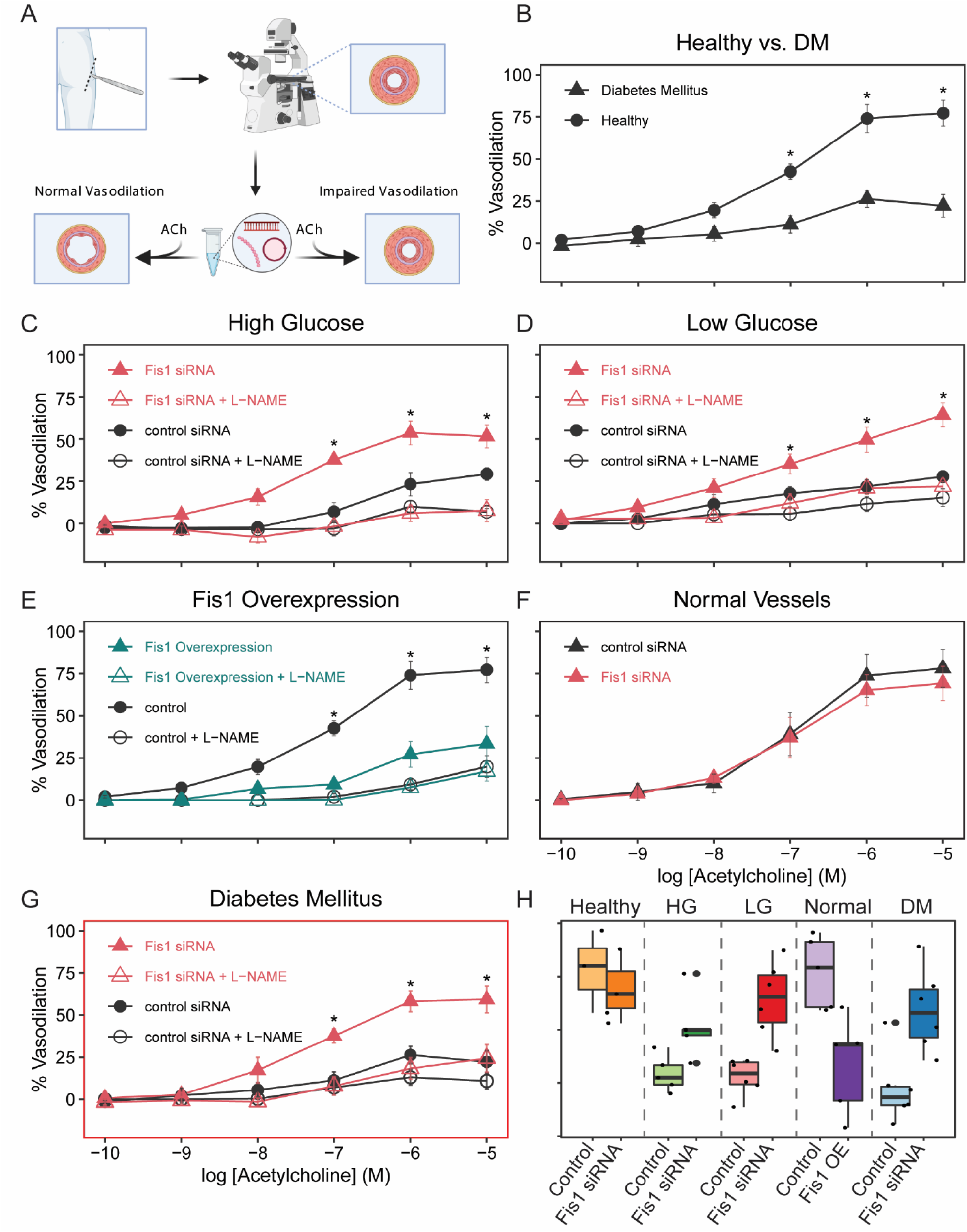
Altered glucose levels impair vasodilation of human resistance vessels in a Fis1-dependent manner. **A.** Schematic of experimental procedures in which human resistance vessels are treated with siRNA or plasmid and vasodilation is measured by videomicroscopy upon acetylcholine treatment. **B.** Vasodilation is impaired in vessels from diabetic patients compared to healthy controls (n = 6 for diabetic vessels and n=5 for healthy vessels, p < 0.0001). Both high **C.** (33 mM, n = 5, 95% CI (-32.4 to -12.1 at 10^-5^ M Ach), p<0.0001) and low **D.** (2.5 mM, n=6, 95% CI (-46.9 to -26.3 at 10^-5^ M Ach), p<0.0001) glucose conditions impair vasodilation which is improved upon Fis1 siRNA treatment in a NO-dependent manner. **E.** Overexpression of Fis1 in healthy vessels significantly attenuates vasodilation (n=5, 95% CI (33.7 to 53.7 at 10^-5^ M Ach), p<0.0001). **F.** Genetic silencing of Fis1 using siRNA does not alter the vasodilatory response of normal vessels (n=3). **G.** Vasodilation of diabetic vessels is significantly improved upon Fis1 siRNA treatment (n=6, 95% CI (-53.9 to -20.2 at 10^-5^ M Ach), p<0.0001). Error bars in vasodilation plots represent SEM. **H.** Maximum vasodilation of vessels exposed to either a control siRNA or Fis1 siRNA in the presence of 10^-5^ M acetylcholine for each of the following conditions: healthy, high glucose, low glucose, and diabetes mellitus, as well as a control treatment or overexpression of Fis1 in healthy vessels. The boxes represent 25^th^ and 75^th^ percentiles, with the horizontal line representing the median % vasodilation.

### Development of peptide inhibitor pep213

In our crystal structure of Fis1 (Dohm *et al*, 2003) (PDB: 1NZN), helix 1 on one protomer binds into a conserved, concave surface of another protomer suggesting that a peptide inhibitor derived from helix 1 might be possible. Indeed, a phage-display screen against a highly truncated Fis1 identified peptides that bound to Fis1 (Serasinghe *et al*, 2010). To confirm binding and determine the manner in which these peptides engaged Fis1, we used uniformly ^15^N-labeled Fis1^1-125^ and performed titration experiments monitored by two-dimensional ^1^H, ^15^N correlation experiments (**Figures 2A and 2E; Supplemental Figures 6A and 6B**). The resulting data were globally fit using TREND (Xu & Doren, 2016) to give apparent *K_D_* values of 171 ± 10 µM and 63 ± 4 µM for pep2 and pep13, respectively (**Figure 2B and 2F**) indicating a higher affinity than previously reported which was likely due to earlier measurements being conducted with an N-terminal His_6_-Myc tag that likely interferes with binding. Given the weak affinities, we sought to create a new peptide with enhanced binding affinity to Fis1. Displaying the NMR chemical shift perturbations from the titration (**Figure 2C and 2G**) onto the Fis1 structure revealed that both peptides bound in a similar manner mediated by similar Fis1 residues that lie on helices 2, 4, and 6 of the concave surface of the molecule (**Figure 2D and 2H**). However, pep2 caused greater perturbation of helix 4 and 6 residues than pep13. These data indicated that a chimera peptide might have higher affinity by incorporating features of both peptides.

**Figure 2.**
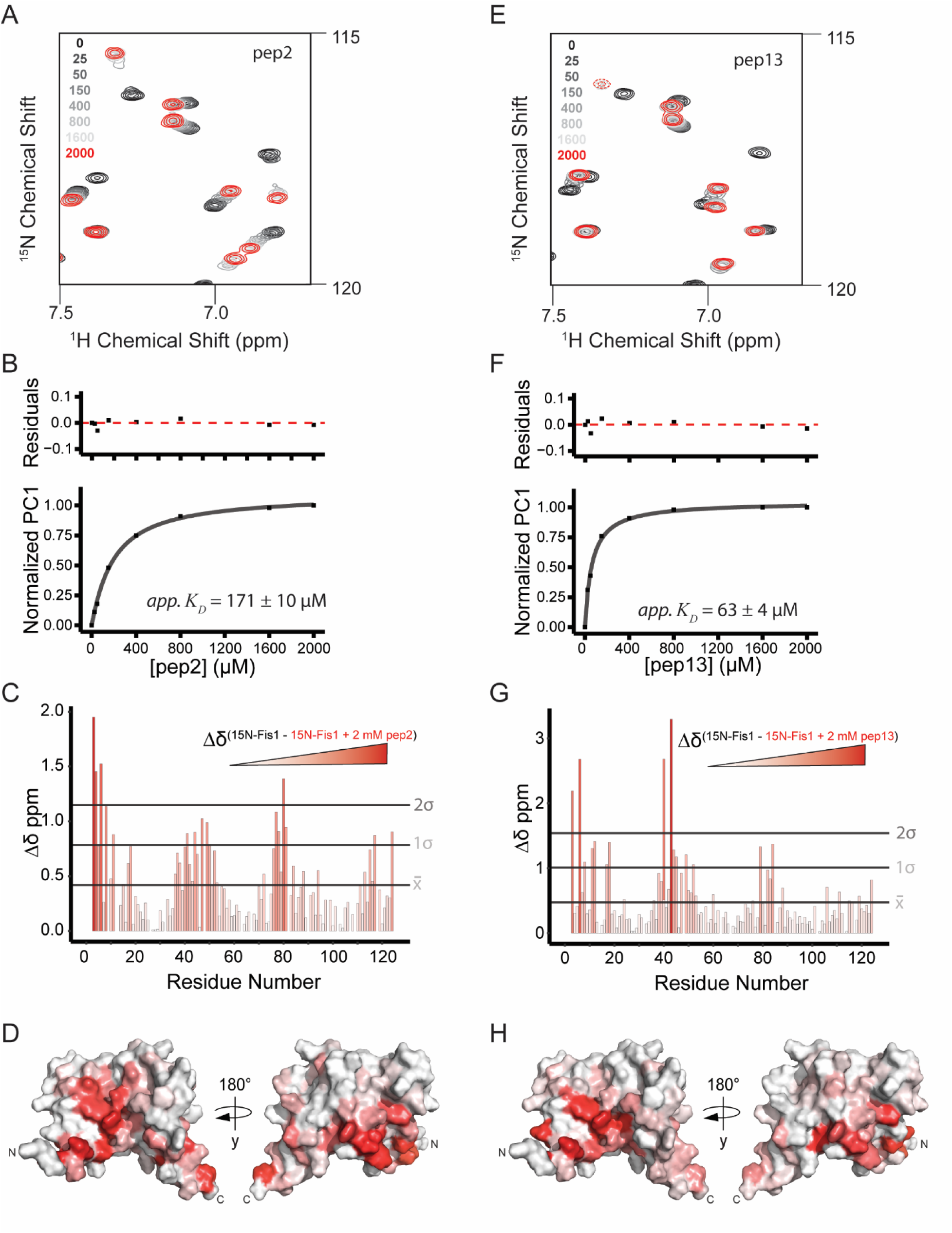
The cytosolic domain of Fis1 binds to pep2 and pep13. **A.** ^1^H, ^15^N HSQC spectral overlays of 50 µM ^15^N-Fis1^1-125^ with increasing amounts (0-2mM) unlabeled pep2 with changes in chemical shift indicative of binding. **B.** Affinity determination of pep2 binding to Fis1 using TREND analysis on all spectra from (**A**) and fitting the resulting normalized PC1 values as a function of pep2 concentration, indicating an apparent *K_D_* = 171 ± 10 µM. **C.** Chemical shift perturbations of Fis1^1-125^ alone or in the presence of 2 mM pep2 (Δδ) are shown for each Fis1 residue in a gradient fashion, where a redder color indicates a greater Δδ. Lines indicate the mean Δδ and both one and two SD from the mean. **D.** Fis1 Δδ values displayed on a surface representation of the structure of Fis1 (PDB: 1PC2^1-125^), in a gradient fashion that replicates the color scheme from (**C**). **E.** ^1^H, ^15^N HSQC spectral overlays of 50 µM ^15^N-Fis1^1-125^ with increasing amounts (0-2mM) unlabeled pep13 with changes in chemical shift indicative of binding. **F.** Affinity determination of pep13 binding to Fis1 as performed in B, indicating an apparent *K_D_* = 63 ± 4 µM. **G.** Chemical shift perturbations of Fis1^1-125^ alone or in the presence of 2 mM pep13 (Δδ) as in (**C**). **H.** Fis1 Δδ values from (**G**) displayed on a surface representation of the structure of Fis1 (PDB: 1PC2^1-125^) as in (**D**). Residuals to the fits shown for **B**, and **F.**

Using NMR-chemical shifts to guide computational modeling, we rationally designed a novel peptide, pep213, that we reasoned might have a higher affinity for Fis1. To evaluate this, we collected 2D NMR experiments on a ^15^N-Fis1^1-125^ sample with increasing amounts of unlabeled pep213 (0-2 mM) (**Figure 3A and 3B**). The NMR chemical shift perturbations from the titration (**Figure 3C**) suggested novel pep213 may bind in a similar location as pep2 and pep13 (**Figure 3D**). TREND analysis indicated an apparent *K_D_* = 7 ± 2 µM (**Figure 3E**), which was confirmed using intrinsic tryptophan fluorescence spectroscopy that gave a similar apparent *K_D_* = 3.3 ± 0.1 µM (**Figure 3F**). To verify peptide sequence specificity in the interaction, a scrambled version of pep213, consisting of the same amino acid composition as pep213 but in random order, showed no chemical shift perturbations upon addition of 2 mM peptide to 50 µM ^15^N-Fis1, indicating the Fis1-pep213 interaction is indeed sequence specific (**Supplemental Figures 6D-E)**.

**Figure 3.**
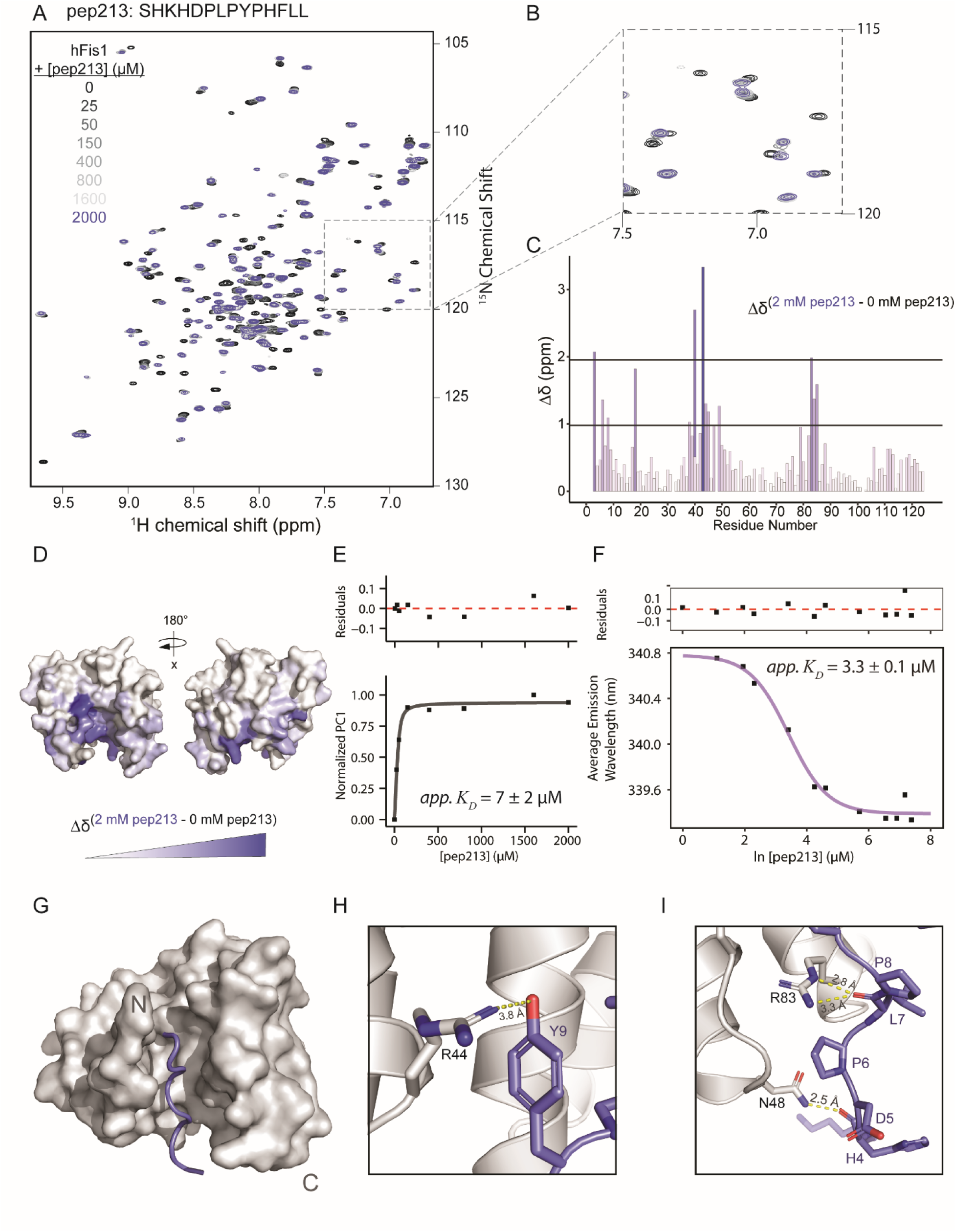
Novel peptide, pep213, specifically binds to the concave surface of human Fis1. (**A**) ^1^H, ^15^N HSQC spectral overlays of 50 µM ^15^N-Fis1^1-125^ with increasing amounts (0-2 mM) unlabeled pep213 with changes in chemical shift indicative of binding. (**B**) Zoomed in region from (**A**). (**C**) The degree of chemical shift perturbation was determined for each residue in ^15^N-Fis1 and then (**D**) mapped onto the structure of Fis1^9-125^ (PDB: 1PC2) with the first eight residues removed for better visualization of the proposed binding site. (**E**) Affinity determination of pep213 binding to Fis1 using TREND analysis on all spectra from (**A**) and fitting the resulting normalized PC1 values as a function of pep213 concentration, indicating an apparent *K_D_* = 7 ± 2 µM. (**F**) Affinity determination of pep213 for Fis1^1-125^ using intrinsic tryptophan fluorescence in which increasing pep213 (0-1 mM) was titrated into 10 µM Fis1, and the resulting average emission wavelength was fit to determine an apparent *K_D_* = 3.3 ± 0.1 µM. Residuals to the fits shown for **E** and **F**. (**G**) Surface representation of Fis1^1-125^ bound to pep213, shown in blue as a ribbon diagram, solved by x-ray crystallography at a resolution of 1.85 Å. Electron density for the two N- and C-terminal residues of pep213 were not visualized. (**H**) Zoomed in region highlights hydrogen bonding between Fis1 R44 and pep213 Y9 and (**I**) Fis1 R83 and N48 with pep213 L7 and H4, respectively.

### Pep213 binds to a highly conserved region of Fis1

The specificity of the pep213-Fis1 interaction was also confirmed by solving a 1.85 Å crystal structure (**Figure 3G, Supplemental Table 1, Supplemental Figure 7**). As predicted from NMR studies, the co-complex structure revealed that pep213 engages the highly conserved Fis1 concave surface (Wells *et al*, 2007), formed by the protein’s tetratricopeptide (TPR) motifs, a common site of protein-ligand interaction among the family of TPR motif-containing proteins (Cortajarena & Regan, 2006; Zeytuni & Zarivach, 2012; Allan & Ratajczak, 2011). The peptide-protein interaction is primarily facilitated through hydrogen bonding and Van der Waals forces.

There is hydrogen bonding between R44 and the peptide Y9 side chain hydroxyl (**Figure 3H**). In addition, there is evidence of hydrogen bonding between the Fis1 N48 residue and peptide H4 backbone carboxyl oxygen, as well as the Fis1 R83 residue with the peptide L7 carboxy oxygen and amide hydrogen (**Figure 3I**). Several of the peptide’s hydrophobic residues, including F7 and L12, face a cleft lined by hydrophobic residues formed by helices α_4_ and α_6_, likely contributing to the low micromolar binding affinity. Interestingly, the peptide’s repeating prolines (PLPYP) allows it to adopt a poly-L-proline type II helix, a third albeit less-common type of secondary structure frequently found at protein-protein interaction interfaces (Adzhubei *et al*, 2013).

Although the two structures were solved in different space groups (1NZN: P6_5_22; co-complex: P4_1_2_1_2) this did not appear to lead to significant structural changes secondary to differential crystal packing. Superimposing the co-complex structure onto the apo structure of 1NZN showed a high degree of similarity, but with an outward displacement of helix α_1_. Given this, it is likely that ligand binding induces conformational changes which are transmitted through the loop between a_1_ and a_2_, as well as a slight shifting of α_2_. Unlike 1NZN, this co-complex structure suggests the N-terminal region of Fis1 is highly disordered and dynamic, more akin to the NMR solution structure (PDB: 1PC2) (Suzuki *et al*, 2003). As such, the N-terminal region appears to almost overlay the peptide and concave surface of Fis1, perhaps facilitated by hydrogen bonding between E7 in the dynamic arm and S45 in a nearby loop. There is not clear evidence of hydrogen bonding between the N-terminal region and the peptide though.

### Pep213 improves endothelial-dependent vasodilation secondary to T2DM, high-glucose exposure, and Fis1 overexpression in human resistance vessels

To test whether pep213 might inhibit Fis1 to improve vasodilation in human arterioles, we first incubated vessels from healthy subjects under high glucose (33 mM) conditions with pep213 fused to the endothelial cell permeant TAT peptide (Brooks *et al*, 2005). Consistent with the siRNA knockdown of Fis1, TATpep213 reversed the poor vasodilatory response in healthy vessels induced by high glucose (**Figure 4A**). Concurrent treatment of TATpep213 with L-NAME abolished the improvements, confirming an NO-dependent effect on vasodilation. This response was specific to pep213 as incubation with TATpep213-scrambled that does not engage Fis1 (**Supplemental Figure 6D-E**) showed no improvement (**Figure 4B**). We next asked if pep213 might improve NO-dependent vasodilation in vessels from T2DM patients, and again we found that TATpep213 induced a significant, and Fis1 specific, recovery of the vasodilatory response of diabetic vessels to Ach (**Figure 4C-D**). To verify pep213’s beneficial effects on vasodilation are indeed due to inhibition of Fis1, we performed a similar experiment in otherwise healthy vessels overexpressing Fis1. In these vessels, we again observed an increase in vasodilation secondary to TATpep213 treatment (**Figure 4E**), albeit with an overall lower maximum vasodilatory capacity compared to healthy vessels under hyperglycemic conditions or T2DM vessels (**Figure 4G**), consistent with pep213 acting specifically through inhibition of Fis1. Importantly, treating normal vessels with TATpep213 did not negatively alter vasodilation compared to the TATpep213-scrambled (**Figure 4F**). These data support that pharmacologically inhibiting Fis1 improves endothelial-dependent vasodilatory function in both diabetic and high glucose-treated vessels.

**Figure 4.**
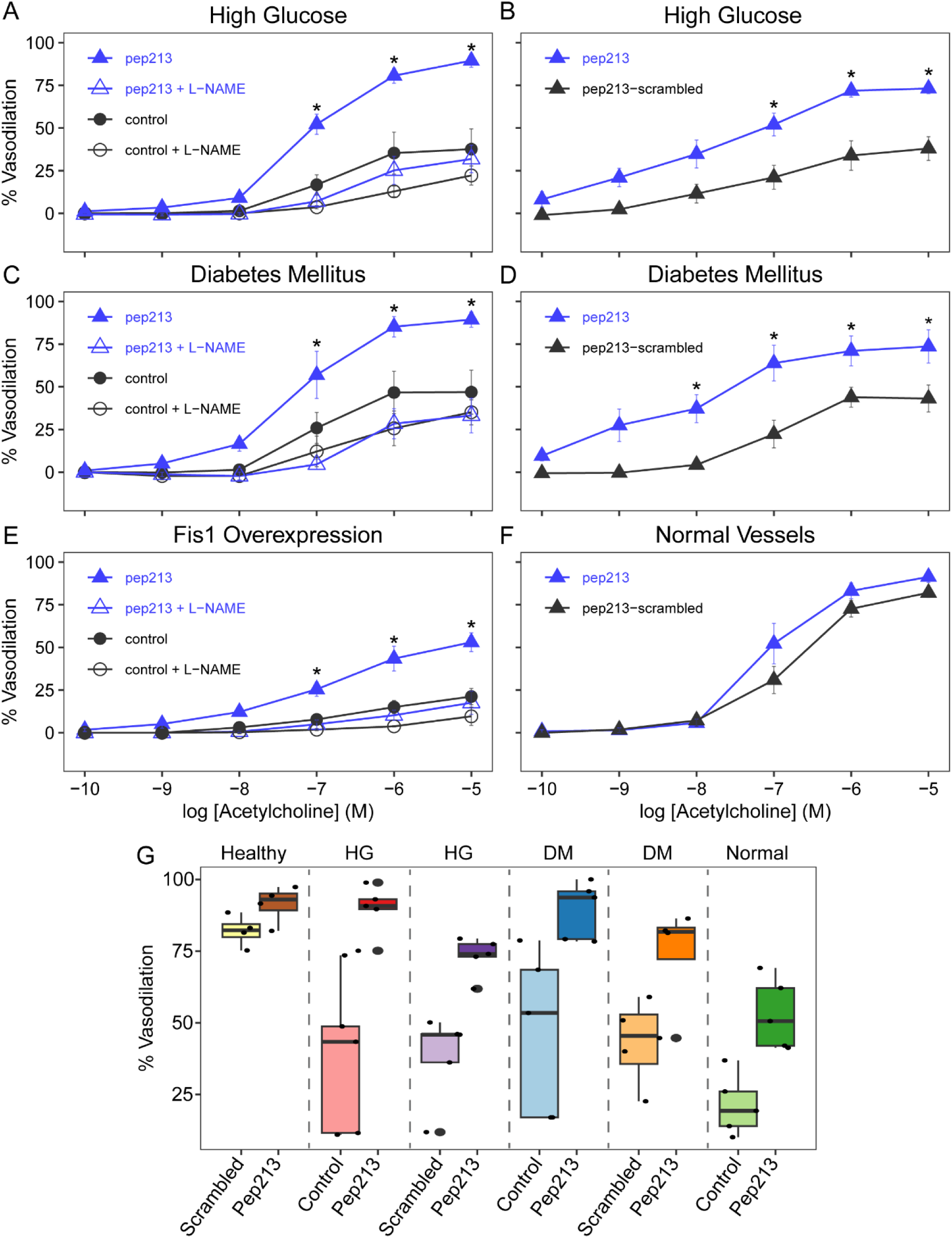
Pep213, but not a scrambled control peptide, reverses impaired endothelium-dependent vasodilation in human vessels from healthy patients. Human resistance vessels isolated from healthy (**A, B**) or T2DM patients (**C, D**) were treated with a TAT-fusion pep213 construct, and vasodilation was measured by videomicroscopy upon acetylcholine treatment. The impaired vasodilation caused by high glucose conditions **A** (33 mM) was reversed by treatment with 1 µM TAT-pep213 compared to control treatment (n=5, 95% CI (-66.9 to -36.9 at 10^-5^ M Ach), p<0.0001) or **B.** compared to a scrambled version of the pep213 sequence fused to a TAT sequence (n=5, p<0.05). **C.** Diabetes-induced impaired vessel vasodilation was rescued upon treatment with 1 µM TAT-pep213 compared to a control treatment (n=5, 95% CI (-57.0 to -14.1 at 10^-5^ M Ach), p<0.001) or **D.** compared to the scrambled peptide control (n=4, p<0.05). **E.** Overexpression of Fis1 in vessels from healthy patients severely attenuates vasodilation which is rescued upon 1 µM TAT-pep213 compared to a control treatment (n=5, 95% CI (-54.8 to -8.7 at 10^-5^ M Ach), p<0.05). **F.** Treatment with 1 µM TAT-pep213 does not alter vessel vasodilatory capacity compared to a scrambled peptide control (n=4). Error bars represent SEM. **G.** Maximum vasodilation of vessels exposed to either TATpep213 and a control siRNA treatment, or TATpep213 and a TATcontrol peptide in which the sequence of pep213 was randomized, in the presence of 10^-5^ M acetylcholine for each of the following conditions: healthy, high glucose, and diabetes mellitus, as well as a control treatment in normal vessels or overexpression of Fis1 in healthy vessels. The boxes represent 25th and 75th percentiles, with the horizontal line representing the median % vasodilation.

### Pep213 treatment reduces mitochondrial-derived O_2_ ^·-^ in vessels from type two diabetic patients and healthy vessels with overexpression of Fis1

It is well established that excess mtROS contributes to the development of endothelial dysfunction in diabetic patients through direct activation of inflammatory pathways such as NF-κB, altered glycolytic activity, and proinflammatory epigenetic changes (Widlansky & Hill, 2018). Therefore, we sought to understand whether inhibition of Fis1 may decrease vessel mtROS levels, hypothesizing that Fis1 may play a driving role in this process given its role in endothelial-dependent impaired vasodilation. To test this, vessels from healthy control patients were treated with a lentiviral Fis1 construct to induce protein expression and the mitochondrial-specific fluorescent ROS indicator MitoNeoD was used to determine levels of mtROS (**Figure 5**). Vessels were then treated with either TATpep213 or TATpep213-scramble for 1 hour. Compared to control vessels which showed minimal fluorescence, vessels overexpressing Fis1 had a significantly higher MitoNeoD fluorescence intensity, suggesting Fis1 overexpression increases ROS production consistent with earlier results in cell-based experiments (Yu *et al*, 2006). When treated with TATpep213, vessel ROS content was significantly decreased (**Figure 5B**). This effect was specific to pep213 as there was no change in mitochondrial ROS upon treatment with the scrambled peptide. Following peptide treatment, the vessels were then incubated with MitoTempo, a ROS scavenging agent, which decreased ROS levels in TATpep213-scrambled vessels to that of the vessels treated with TATpep213. However, there was no further improvement in ROS levels in the TATpep213 treated vessels, suggesting mtROS was already improved by pep213, and that mechanistically, excess Fis1 likely induces endothelial dysfunction and resultant impaired vasodilation through increased generation of mtROS.

**Figure 5.**
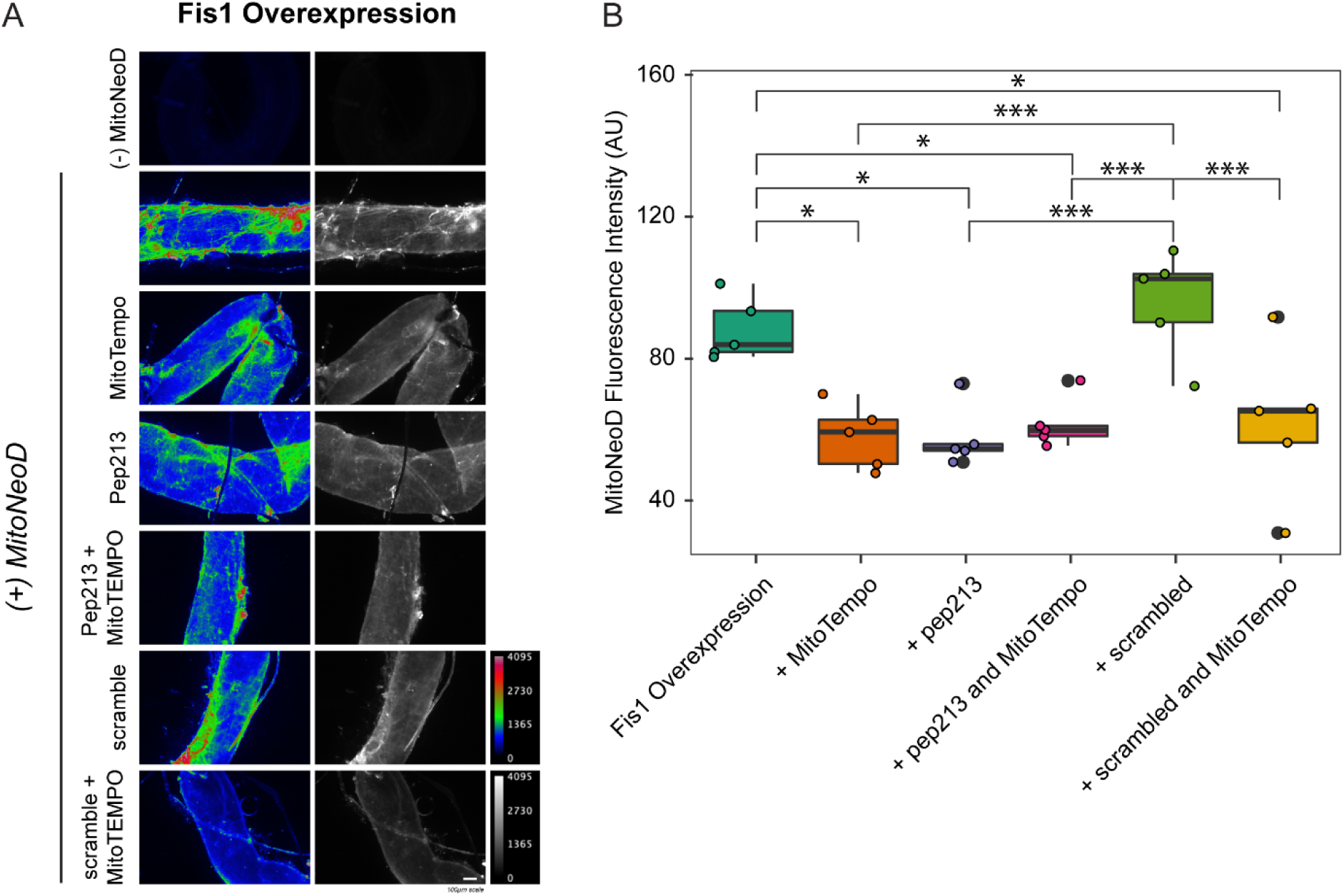
Pep213 treatment reduces mitochondrial-derived O2^·-^ in vessels overexpressing Fis1. **A.** Representative images of vessels overexpressing Fis1 were stained with the mitochondrial-specific superoxide probe MitoNeoD and treated with either TATpep213, a TATscrambled-pep213 for control, the ROS scavenging agent MitoTEMPO, or a combination of peptide and MitoTEMPO. Fluorescence intensity of MitoNeoD, pseudocolored in a rainbow gradient where red corresponds to increased ROS. **B**. Quantification of MitoNeoD fluorescence intensity in vessels indicates decreased ROS upon pep213 treatment, which is not further improved upon co-treatment with MitoTEMPO, but no improvements in ROS with the scrambled peptide control (n=5, *p<0.05, ***p<0.001). The boxes represent 25^th^ and 75^th^ percentiles, with the horizontal line representing the median fluorescence intensity.

## Discussion

The present studies implicate Fis1 as having a critical role in diabetic endothelial dysfunction in human vessels and support pharmacologically targeting Fis1 to improve vascular health in patients with diabetes. Consistent with this are our findings that molecular suppression of Fis1 or inhibition with a first-in-class Fis1 inhibiting agent, pep213, improves impaired NO production and vasodilation in both diabetic vessels and healthy vessels exposed to abnormal glucose conditions. These favorable effects occur without changes in endothelial cell metabolism or altered expression of other proteins involved in mitochondrial fusion, fission, or autophagy. Importantly, we found that both high and low glucose conditions, in addition to T2DM, induce this impaired vasodilation in a Fis1-dependent manner, suggesting these effects are likely glucose-driven as opposed to deriving from the multifactorial effects of diabetes mellitus.

Through the solution of a co-complex structure, we found that pep213 binds a concave surface on Fis1 within its highly conserved TPR region. In yeast Fis1, this interface is the site of interaction with the cytosolic mitochondrial fission protein Dnm1lp (yeast Drp1) (Wells *et al*, 2007). Therefore, the beneficial actions of pep213 may stem from the inhibition of a protein-protein interaction between Fis1 and Drp1. Although it is well established that yeast Fis1 is directly involved in yeast mitochondrial fission, the role of Fis1 in human mitochondrial fission remains unclear. Given the evolution of other Drp1 recruiting proteins, many in the field suggest Fis1 has adopted other cellular roles, such as mitophagy (Ihenacho *et al*, 2021), likely through interactions with the Rab7 and Rab8 modulators TBC1D15 and TBC1D17, as well as Syntaxin17(Onoue *et al*, 2013; Iwasawa *et al*, 2011; Yamano *et al*, 2014; Xian *et al*, 2019). In addition, recent work has suggested Fis1 may act in fission through inhibition of the mitochondrial fusion proteins Mfn1, Mfn2, and OPA1 (Yu *et al*, 2019), as opposed to direct Drp1 recruitment. Emerging data suggests Mfn2 expression levels may regulate mitochondrial membrane potential, mitochondrial ROS generation, and NO bioavailability (Bach *et al*, 2003; Lugus *et al*, 2011). Given Mfn2 has also been shown to be downregulated in the endothelium of diabetic patients (Shenouda *et al*, 2011), the role of mitochondrial fusion in the regulation of human vascular endothelial function merits additional investigation. However, recent work suggests that Fis1 does appear to be directly involved in mediating peripheral fission events whereas Mff, the primary Drp1 recruiter (Otera *et al*, 2010), mediates central fission events (Kleele *et al*, 2021), suggesting divergent roles in Drp1 recruitment.

It is possible that Fis1 may mediate only certain types of mitochondrial fission, such as under high cellular stress induced by hypoxia and abnormal glucose conditions (Kim *et al*, 2011; Kumari *et al*, 2012; Ciarlo *et al*, 2010; Kaddour-Djebbar *et al*, 2010). Supporting this, Fis1 has shown to be over-expressed in endothelial cells in the setting of diabetes, as well as during acute exposure to high glucose conditions or excessive free fatty acids (Shenouda *et al*, 2011; Liu *et al*, 2017; Gioscia-Ryan *et al*, 2016; Choi *et al*, 2016; Zhang *et al*, 2018; Yu *et al*, 2006, 2008). Further, molecular silencing of Fis1, or Drp1, in aortic endothelial cells exposed to high glucose results in an increase in phosphorylation of eNOS at its Ser1177 activation site, allowing for increased NO production and resultant vasodilation (Shenouda *et al*, 2011), consistent with our results here. A primary role in stress-induced fission makes Fis1 a promising therapeutic target as its inhibition may still allow for homeostatic mitochondrial fission to occur.

Properly maintained mitochondrial networks are critical for supporting normal mitochondrial function and allows for organellar distribution and maintenance to meet cellular metabolic demands, removal of damaged mitochondria, and limiting of mtROS production (Chen & Chan, 2009; Chan, 2012; Detmer & Chan, 2007; Suen *et al*, 2008; Yu *et al*, 2006). Long-term exposure to excess glucose, such as in diabetes mellitus, stimulates numerous metabolic pathways including the TCA cycle. This results in an overproduction of NADH and FADH_2_ with resultant increases in the mitochondrial proton gradient and superoxide production (Brownlee, 2001). Excess nutrients, as well as starvation conditions, also stimulate mitochondrial networks to undergo fission in multiple cell types, including endothelial cells, in conjunction with the increased mitochondrial ROS production (Shenouda *et al*, 2011; Yu *et al*, 2006; Molina *et al*, 2009; Jheng *et al*, 2012; Tanner *et al*, 2017).

Previous work has shown that mitochondrial fragmentation occurs upstream of ROS generation and increased cellular respiration rates secondary to hyperglycemic conditions (Yu *et al*, 2008; Galloway *et al*, 2012). Although inhibition of mitochondrial fission using an enzymatically inactive Drp1 K38A variant prevents increased mtROS generation under high glucose settings, the reverse is not true, and inhibition of mtROS generation does not prevent hyperglycemic induced fission (Yu *et al*, 2008). Conversely, the opposite effect has been observed in models of ischemia-reperfusion injury, in which mtROS appears to drive mitochondrial fission and can be rescued by treating with antioxidants (Giedt *et al*, 2012), suggesting the instigating factor in this cycle may be contextually dependent. Our results show that Fis1 inhibition, and therefore presumably inhibition of either mitochondrial fission or mitophagy, decreases mtROS production, suggesting, at least under hyperglycemic conditions, it is excess mitochondrial fission driving mtROS. Seminal work performed in cultured endothelial cells demonstrated that high glucose-driven mtROS promotes both acute and chronic impairment in function through numerous cell signaling and epigenomic pathways (El-Osta *et al*, 2008; Nishikawa *et al*, 2000). Strikingly, by directly reducing mitochondrial superoxide levels in resistance arteries from T2DM patients, impaired endothelium-dependent vasodilation can be reversed, emphasizing this important connection between mtROS and endothelial function (Kizhakekuttu *et al*, 2012). The decrease in vessel mtROS levels upon pep213 treatment in our studies suggests this may be one way by which pharmacological inhibition of Fis1 improves endothelial dysfunction.

Our results suggest that endothelial dysfunction is an NO-dependent phenomenon and does not derive from altered cellular bioenergetics. It is perhaps surprising that neither exposure to high glucose conditions nor Fis1 knockdown had any change on cellular bioenergetics given previous reports of increased oxygen consumption rates under high glucose conditions (Galloway *et al*, 2012; Yu *et al*, 2006). However, ATP synthesis rates are dictated primarily by ATP demand, which may not be altered under basal conditions, as well as to a lesser extent, nutrient availability, and intrinsic proton leak (HAFNER *et al*, 1990). In addition, GLUT1 is considered the primary glucose transporter in endothelial cells and has a *K_M_* for glucose between 3.4-20 mM (Palfreyman *et al*, 1992; Lundqvist & Lundahl, 1997; Kasahara & Kasahara, 1997; KASAHARA & KASAHARA, 1996). Therefore, the GLUT1 transporters would likely become saturated with even modest increases in glucose concentrations and an increase in glucose to 33 mM, as was used in this study, would not be expected to significantly alter intracellular glucose concentrations or bioenergetics. Our results may also be representative of cell-type specific effects or abnormal transporter localization due to a loss of cellular polarity as has been observed previously in cultured human coronary artery endothelial cells (HCAECs) (Gaudreault *et al*, 2008).

Curiously, several of the medications used for managing T2DM also inhibit mitochondrial fission as an “off-target” effect by reducing Fis1 and/or Drp1 expression. Empagliflozin, an SGLT2 inhibitor with favorable effects on cardiovascular risk, mortality, and microvascular renal disease in T2DM patients (Zinman *et al*, 2015; Wanner *et al*, 2016), decreases both Fis1 over-expression and mitochondrial fission, and up-regulates mitochondrial superoxide dismutase (SOD) in rat cardiac myocytes (Mizuno *et al*, 2018). Metformin, a long-time first-line agent for T2DM glucose control with favorable cardiovascular effects, reduces atherosclerotic formation in ApoE^-/-^ mice by suppressing Drp1-mediated fission (Wang *et al*, 2017). The dipeptidyl peptidase 4 inhibitor, vildagliptin, from a class of medications known to increase NO production, reduces expression of Fis1 and Drp1, Drp1 translocation to mitochondria from the cytosol, and both mitochondrial fission and ROS production, while increasing NO production in the aortic endothelium of diabetic mice (Liu *et al*, 2019). Given our results, we predict these common anti-diabetes medications may have ameliorative vascular effects in T2DM in part based on off-target effects on the expression and/or interaction of Fis1 and Drp1. Whether these improvements are due solely to improved glycemic control, or perhaps direct inhibition of Fis1 or Drp1, merits future study.

In this paper, we present a first-in-class Fis1 inhibitor, pep213, with promising therapeutic potential for both T2DM- and high glucose-induced impairments of endothelium-dependent vasodilation due to excess ROS generation. Recently, Fis1 has also been implicated in the pathogenesis and progression of numerous neurodegenerative disorders including Alzheimer’s disease, amyotrophic lateral sclerosis (ALS), Huntington’s disease, and septic encephalopathy (Haileselassie *et al*, 2020; Joshi *et al*, 2018b; Guo *et al*, 2013; Joshi *et al*, 2018a; Qi *et al*, 2013), as well as septic cardiomyopathy and ischemia-reperfusion injury post myocardial infarction (Disatnik *et al*, 2013; Haileselassie *et al*, 2019). Initial efforts to inhibit Drp1 have shown great promise in models of these diseases, and we predict that direct pharmacological inhibition of Fis1 may prove beneficial for treating or preventing the progression of these pathologies as well.

## Materials and Methods

### Subject recruitment and screening

Human adipose resistance arterioles were obtained using two research protocols that were reviewed and approved by MCW’s Institutional Research Board. We recruited 67 individuals with T2DM and healthy individuals (ages 21-75;14 T2DM and 53 healthy controls) without cardiovascular risk factors as previously described for gluteal adipose pad biopsies (Kizhakekuttu *et al*, 2012; Widlansky *et al*, 2018). Individuals with T2DM were either on medications to treat T2DM and/or met criteria for the diagnosis of T2DM as described by the American Diabetes Association (Association, 2018). Study methods were reviewed and approved by the Institutional Research Board of the Medical College of Wisconsin and all subjects signed a written informed consent form prior to proceeding with any study activities. All experiments conformed to the principles outlined in the WMA Declaration of Helsinki and the Department of Health and Human Services Belmont. All subjects were screened to assure they met study inclusion criteria. In all subjects, height and weight were measured once, and heart rates and blood pressures measured in triplicate. Subjects were excluded if they had any of the following: known atherosclerotic disease (coronary artery disease, peripheral vascular disease, history of stroke or myocardial infarction), chronic liver disease, elevated plasma creatinine (> 1.5 mg/dL in men, > 1.4 mg/dL in women), a diagnosis of cancer in remission less than a year, regularly taking a blood thinner or anti-platelet agent other than aspirin, or smoking cigarettes within a year of enrollment. Individuals without T2DM were also excluded if they had an LDL cholesterol ≥ 160 mg/dL, hypertension (blood pressure ≥ 140/90 mmHg), or if they were on medications to treat either of these conditions. Vessels from an additional 18 subjects (14 non-DM, 4 T2DM) were obtained from discarded subcutaneous adipose tissue. The demographics and available clinical information for subjects samples used in each experiment are included in **Supplemental Tables 2-8.**

### Human resistance artery acquisition by gluteal adipose pad biopsy

For 67 subjects, human resistance arteries from adipose samples were obtained from the upper outer quadrant of the gluteal adipose pad as previously described (Kizhakekuttu *et al*, 2012; Tanner *et al*, 2017; Widlansky *et al*, 2018; Suboc *et al*, 2013; Mohandas *et al*, 2015). Briefly, following sterilization and anesthesia with 1% lidocaine, a 1-1.5 cm incision was made in the upper outer quadrant of the gluteal adipose pad. An approximately 8 cm^3^ volume of adipose tissue was then removed by sharp dissection. After achieving hemostasis, the wounds were closed with 1-2 absorbable deep dermal sutures and the epidermal layer was closed using either Dermabond or Steri-Strips.

### Cell culture

Immortalized human microvascular endothelial cells (HMEC-1) purchased from ATCC (Manassas, VA) were cultured in antibiotic-free MCDB131 (Life Technologies, Carlsbad, CA) supplemented with 10 mM Glutamine (Thermo Fisher Scientific, Waltham, MA), 10% FBS (Sigma-Aldrich, St. Louis, MO), 10 ng/mL Human EGF (Thermo Fisher Scientific), and 1 µg/mL Hydrocortisone (Sigma-Aldrich). For assays involving high glucose (33 mM), normal glucose (5 mM), or low glucose (2.5 mM) conditions, antibiotic-free MCDB131 was supplemented as above and glucose levels were adjusted by adding glucose in sterile PBS. Human dermal microvascular endothelial cells (HMEC) were purchased from Lonza (Basel, Switzerland) and supplemented with Microvascular Endothelial Cell Growth Medium-2 Bullet Kit (Lonza).

### Transfection protocols for cultured endothelial cells and human vessels

#### Transfection of Fis1 and Drp1 siRNA and scrambled control into cultured cells

Constructs for Fis1 and Drp1 siRNAs were acquired from Origene (Rockville, MD, sequences included in the supplemental material). Lipofectamine RNAi Max (Thermo Fisher Scientific) vehicle was added to 20 nM of RNAi constructs in Opti-MEM (Life Technologies). The mixture was diluted in culture media and incubated with HMEC-1 cells for four hours prior to replacement with fresh culture media (MCDB131). Cells were subsequently incubated for 24 hours before further treatment and analyses. Following incubation, cells were treated with either high glucose for 6 hours, or low glucose for 2 hours, prior to measuring NO production, bioenergetics, protein expression, and endothelial layer integrity.

#### Transfection of Fis1 and Drp1 siRNA and scrambled controls in human resistance arteries

Transfection of human resistance arteries with siRNA constructs was performed as previously described (Widlansky *et al*, 2018; Tanner *et al*, 2017). Resistance arteries dissected from adipose tissue were suspended in a culture myograph chamber (204 CM, DMT, Ann Arbor). One end of the vessel was sutured onto a glass pipette within a micro-organ chamber. Prior to suturing the second end of the arteriole to a glass pipette, Fis1 siRNA, Drp1 siRNA (20 nM in optiMEM using Lipofectamine RNAiMax, Invitrogen) or control scrambled siRNA (20 nM) were gently injected into the vessel lumen. The loose end of the vessel was then tied onto another glass pipette, and the vessel was suspended in the chamber and placed on a myograph stage bathed in Krebs buffer at 37°C and pressurized to 60 mmHg. Following 4-6 hours of incubation, the siRNA was slowly washed out of the lumen over a 24-hour period at a low shear rate (< 5 dyn/cm^2^).

#### Transfection for Fis1 Overexpression in Human Vessels

Plasmid CDH5-AbR-GFP (Addgene Cat# 122970) containing endothelial specific-promoter (CDH5, VE-Cadherin) and green fluorescent protein (GFP) and the plasmid pC4-RhE-FRB-Fis1 (Addgene Cat# 68056) containing human Fis1 gene were purchased from Addgene. The working viral vectors were constructed from these two plasmids by Custom DNA Constructs, LLC (Islandia, NY). Lentiviral particles (pL6-CDH5p-EGFP and pL6-CDH5p-GFP-T2A-Fis1) were constructed in the viral core lab at the Versiti Blood Institute (Milwaukee, WI). Lentiviral particles pL6-CDH5p-EGFP or pL6-CDH5p-GFP-T2A-Fis1 were gently injected into the lumen of human arterioles at concentration of 50 transfection units (TU)/cell. Following injection of each viral preparation into the vessel lumen, both ends of the vessel were secured with 10-0 nylon monofilament suture (22 µm diameter). Vessels were kept in culture media containing Penicillin-Streptomycin and incubated at 37°C, 5% CO_2_, and > 95% humidity for 48 hours. Over-expression for Fis1 and endothelial specificity was verified by fluorescent microscopy (Ex./Em. 488/535 nm).

### Wes for quantification of protein knock-down in human resistance arteries

Microvessels were rinsed twice with cold Cell Wash Buffer (ProteinSimple, San Jose, CA) and the vessels were placed in 0.5 mL tubes containing 1.4 mm ceramic beads (Omni International, Kennesaw, GA) containing 35 µl cold RIPA lysis buffer (NanoPro® Cell Lysis Kit, ProteinSimple, San Jose, CA) for tissue homogenization on the Bead Ruptor 12 (Omni International, Kennesaw, GA). The vessels were homogenized under the following conditions: speed: 6.0 m/s, time: 20 secs, cycles: 3, dwell time between cycles: 3. Samples underwent a quick spin and lysates were transferred to new 1.7 mL microcentrifuge tubes. Lysates were centrifuged at 14,000 x *g* (Eppendorf 5424 R, Hamburg, Germany) for 15 minutes and flash frozen in liquid nitrogen and subsequently stored at -80°C. Protein concentration was quantified by the *DC*™ Protein Assay (Bio-Rad, Hercules, CA) to ensure loading of equal total protein amounts and the subsequent protein expression was evaluated by the automated capillary electrophoresis-based immunodetection via Simple Western system, Wes™ (ProteinSimple, San Jose, CA). Proteins were analyzed and detected with the 12-230 kDa Wes™ separation module with a 25-capillary cartridge. Lysates (run in duplicate), blocking buffer, a 1:20 dilution of Fis1 primary antibody (#104956-1-AP, ProteinTech, Rosemont, IL), anti-Rabbit secondary antibody (Bio-Techne, Cat #042-206, undiluted), chemiluminescent substrate, a biotinylated size marker, total protein detection reagents, and wash buffer were loaded in the designated wells on the supplied microplate according to manufacturer’s instructions. The plate was centrifuged for 5 min at 1000 x *g* then loaded into the Wes™ instrument set to default separation parameters. Compass for Simple Western software (version 3.1.7, ProteinSimple) was used for data analysis and integration of specific antibody peaks. Protein expression was normalized to total protein which was determined via an additional Wes assay with biotin-based protein labeling and analyzed using Compass for Simple Western software as the integral of the antibody peaks. Each measure was normalized to sample total protein level.

### Measurement of endothelium-dependent vasoactivity

Transfected resistance arteries from healthy individuals were exposed to either normal glucose conditions (5 mM), 2 hours of low glucose (2.5 mM) or 6 hours of high glucose (33mM). All studies in vessels from patients with T2DM were performed under normal glucose conditions, whereas healthy control vessels were used for both the low and high glucose studies. Vessels were pre-constricted to approximately 50% of their resting diameter with endothelin-1 (Sigma Aldrich, USA). The constricted arterioles were then exposed to acetylcholine (Sigma Aldrich, USA) at serially increasing concentrations from 10^-10^ to 10^-5^ M and changes in vessel diameter were measured using digital calipers and videomicroscopy. After the final exposure to the 10^-5^ M dose of acetylcholine, vessels were exposed to 100 µM papaverine to test their smooth muscle reactivity. Following a 10-minute washout period, the same arterioles were incubated with *N_ω_*-Nitro-L-arginine methyl ester (L-NAME, 100∝M), a direct inhibitor of nitric oxide synthase, for 30 minutes and vasodilation was subsequently remeasured following pre-constriction by ET-1 and exposure to increasing concentrations of acetylcholine (10^-10^ to 10^-5^ M). For studies using pep213, 1 ∝mol/L of pep213 attached to a TAT-sequence or scrambled peptide control attached to a TAT-sequence were injected into the lumen of vessels and incubated for 1 hour in an organ chamber with Kreb’s buffer continuously bubbled with gas mixture at 37°C prior to vasoactivity measurements.

### Measurements of Nitric Oxide (NO) Bioavailability

#### Cells in culture

To measure nitric oxide (NO) bioavailability in HMEC-1 cells and arterioles, a fluorescent NO marker, 4,5-diaminofluorescein diacetate (DAF-2 DA, Cayman Chemical) was used. In a dark room, cells were incubated with or without 100 µM L-NAME for 2 hours followed by 15 minutes of incubation with 5 µM DAF-2 DA at 37^○^C. Fluorescence intensity was measured using a SPECTRAFluor Plus plate reader (Tecan, Morrisville, NC) using excitation and emission wavelengths of 485 nm and 535 nm, respectively.

#### Human resistance arteries

For these experiments, two arterioles were isolated from each subject and each vessel cut in half to facilitate four different exposures for each subjects’ vessels. Exposure states included transfection with Fis1 siRNA, Drp1 siRNA, or scrambled control siRNA with and without concomitant exposure to L-NAME at room temperature for 30 minutes. Vessels were either exposed to LG (2.5 mM for 2 hours), HG (30mM for 6 hours), or maintained with exposure to normal glucose concentrations (5 mM) prior to exposure to acetylcholine or DAF2-DA. A subset of vessels were exposed to L-NAME (100 µM) just prior to incubation with DAF2-DA. Acetylcholine (10^-5^ M) and DAF-2 DA (5 µM final concentration) were subsequently added to each vessel and incubated at 37°C for 30 minutes. The arterioles were then washed with PBS buffer, mounted on a slide, and imaged by fluorescence microscopy. Untreated and unstained arterioles were used as controls for fluorescence background signal. Both the treated and control vessels were measured using the same gain settings and the results were analyzed using Meta Morph 7.8 software with statistical significance determined using Friedman’s test with post-hoc Dunnett’s multiple comparison analysis (Universal imaging, West Chester, PA).

### Measuring Mitochondrial Superoxide Levels in Human Resistance Arterioles

Vessels were incubated with 10 µM of MitoNeoD (Medkoo Biosciences, Morrisville, NC) in PBS for 20 minutes in a 37^°^C water bath. Vessels were washed three times with PBS prior to measuring fluorescence intensity. As a control to measure non-specific oxidation, we concomitantly incubated half of the vessels with MitoTEMPO (100 µM). Fluoresence intensity was measured by fluorescence microscopy with Ex./Em. of 594/615 nm (Red). The fluorescence intensity was analyzed by using MetaMorph 7.8.6 software (Universal Imaging, West Chester, PA).

### Measurement of endothelial layer integrity

Human microvascular endothelial (HMEC-1) cells were seeded at 40,000 cells/well and grown on gold electrode array plates (8W10E+, Applied Biophysics Inc.) until approximately 50% confluent. The cells were pre-transfected with Fis1 siRNA (20 nM) or scrambled siRNA (20 nM). On the days of experiments, the transfected cells were exposed to different glucose conditions: high glucose (33 mM) for 6 hours, normal glucose (5 mM) for 2 hours, or low glucose (2.5 mM) for 2 hours. The integrity of the monolayers was checked at 64,000 Hz with less than 10 nF capacitance. The cells were then subjected to an electric cell-substrate impedance sensing (ECIS) functional assay, and monolayer transepithelial electrical resistance (TEER) was measured in real-time using an ECIS ZTheta Instrument (Applied Biophysics Inc., Troy, NY). Resistance was measured over the following frequencies, with 4,000 Hz as the standard: 125 Hz, 250 Hz, 500 Hz, 1,000 Hz, 2,000 Hz, 4,000 Hz, 8,000 Hz, 16,000 Hz, 32,000 Hz, and 64,000 Hz.

### Mitochondrial bioenergetic measurements

HMEC-1 cells were seeded (20,000 cells/well) onto a 96-well Seahorse microplate 4-6 hours before treatment to let the cells adhere to the plate. Cells were pre-transfected with siRNA targeting Fis1 or a siRNA scrambled control. The medium was aspirated and replaced with either medium containing high glucose (33 mM) for 6 hours, or normal (5 mM) or low (2.5 mM) glucose for 2 hours. The high, normal, or low glucose medium was then removed and replaced with XF Base Medium Minimal Dulbecco’s modified Eagle’s medium (DMEM, pH 7.4) supplemented with glucose 45%, L-Glutamine (200 mM), and sodium pyruvate (100 mM) with the final glucose concentration the same as the initial medium concentration. Cells were then incubated in a CO_2_-free incubator at 37°C for 1 hour. During mitochondrial stress tests, the following drugs were sequentially injected in increments of 25 μL: Oligomycin A (2.5 μM final well concentration), FCCP (carbonyl cyanide 4-(trifluoromethoxy) phenylhydrazone, 1 μM final well concentration), and rotenone & antimycin A (1 μM final well concentration each). During glycolytic stress tests the following substances were sequentially injected in increments of 25 μL: glucose (10 mM final well concentration), oligomycin A (2.5 μM final well concentration), and 2-deoxyglucose (50 mM final well concentration). Oxygen consumption rates (OCR) and extracellular acidification rates (ECAR) were then measured using a XFe96 analyzer (Seahorse Biosciences, North Billerica, MA, USA. Following these measurements, cells were fixed with 4% paraformaldehyde for 15 minutes and stained with DAPI (4’,6-diamidino-2-phenylindole, 1:500 dilution of of 1.5 μg/mL stock) for 24 hours. Cell counts were obtained using automated cell counting on a Cytation5 cell imaging multi-mode reader (BioTek, Vermont, USA). OCR and ECAR measurements were then normalized to cell number.

### Measurement of mitochondrial proteins

HMEC-1 cells were grown in six well plates at 0.3 X 10^6^ cells/well and transfected with either siFis1 or a control siRNA. After transfecting for 4-6 hours, the medium was changed to remove exposure to further siRNA and incubated overnight. Cells were then treated with media containing either high glucose (33 mM) for 6 hours, or normal (5 mM) or low (2.5 mM) glucose for 2 hours. The plates were then washed twice with cell wash buffer and 50 µl RIPA lysis buffer (ProteinSimple, San Jose, CA) was added to each well. The cells were scraped gently on ice and transferred to a labeled tube. Lysates were centrifuged at 10,000 RPM (9,400 x g) (Eppendorf 5424 R, Hamburg, Germany) for 10 minutes and flash frozen in liquid nitrogen and subsequently stored at -80°C overnight. The cell pellet and supernatant were thawed, vortexed briefly, and centrifuged at 10,000 RPM (9,400 x g) for 10 minutes. The supernatant was transferred to labeled tubes, protein concentration was quantified by Bradford assay to ensure loading of equal total protein amounts, and protein expression was evaluated by automated capillary electrophoresis-based immunodetection via WES (ProteinSimple). Proteins were analyzed and detected with the 12-230 kDa WES separation module with 25 capillary cartridges. Lysates were first diluted in 0.1X sample buffer in a 4:1 ratio of sample to fluorescent master mix (ProteinSimple), then denatured by heating at 95°C for 5 minutes. For immunodetection by WES, blocking buffer, primary antibody (α-Fis1: Proteintech 10956-1-AP rabbit polyclonal, 1:10 dilution), secondary antibody (α-rabbit secondary HRP: Bio-Techne, Cat #042-206, undiluted), chemiluminescent substrate, the denatured samples, a biotinylated size marker, and wash buffer were loaded in the designated wells on the supplied microplate. The plate was centrifuged for 5 min at 1000 x g then loaded into the WES instrument. Protein detection used default separation parameters. Compass for Simple Western software (version 3.1.7, ProteinSimple) was used for data analysis and integration of specific antibody peaks. Protein expression was normalized to total protein which was determined via an additional WES assay with biotin-based protein labeling and analyzed using Compass for Simple Western software as the integral of the antibody peaks. Control carryover samples between plates were used to correct for inter-plate signal intensity variability. Each measure was normalized to total sample protein level.

### Synthesis of peptides

Peptides pep2 (SHKQDPLPWPRF), pep13 (KHDPLPYPHFLL), pep213 (SHKHDPLPYPHFLL), TAT-pep213 (YGRKKRRQRRRGSGSGSSHKHDPLPYPHFLL), and a TAT-scrambled pep213 (YGRKKRRQRRRGSGSGSPLLHFHPSHDLYPK) were all purchased from Genscript (Piscataway, NJ) with N-terminal acetylation and C-terminal amidation which were determined to be >95% purity by HPLC. The TAT-pep213 fusion peptide included a GSGSGS linker between the TAT cell penetrating sequence (YGRKKRRQRRR) and pep213.

### NMR titration experiments

To determine peptide binding affinities for Fis1, NMR titration experiments were performed similarly to chemical fragment titrations as previously described^17^. First, peptides were resuspended in Fis1 dialysate buffer (100 mM HEPES pH 7.4, 200 mM NaCl, 1 mM DTT, 0.02% v/v sodium azide) to a final concentration of 6 mM. Then, 220 µL of 50 µM ^15^N–hFis1 and increasing concentrations of peptide (0, 25, 50, 150, 400, 800, 1600, and 2000 µM) were prepared and loaded into 3 mm NMR tubes. For each sample, ^1^H, ^15^N HSQC spectra were collected at 25 °C on a Bruker Avance II 600 MHz spectrometer equipped with a triple resonance z-axis gradient cryoprobe and SampleJet autosampler, which allowed automatic tuning, shimming, and data collection for each sample. ^1^H, ^15^N HSQC experiments consisted of 8 scans with 1024 and 300 complex points in the ^1^H and ^15^N dimensions, respectively. Spectra were processed with automated python scripts using NMRPipe and chemical shifts were measured using TitrView and CARA software (Delaglio *et al*, 1995; Masse & Keller, 2005). Peptide binding affinity was determined by TREND analysis to each spectrum within the titration series (Xu *et al*, 2017; Xu & Doren, 2016). TREND reveals the changes in data by performing principal component analysis, where each concentration point is treated as a unique data point; here, the input is each ^1^H, ^15^N spectrum with increasing amounts of peptide. After performing principal component analysis, the principal component 1 (PC1) values are normalized to the highest PC1 value resulting in a range from 0 to 1. Then, the normalized PC1 values are plotted against peptide concentration and fit to a ligand depletion function with protein concentration held constant (Equation 1). Spectral overlays were generated using XEASY software and Adobe Illustrator.

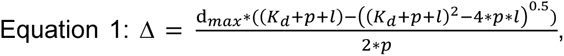

where Δ = adjusted chemical shift change, d_max_ = maximum chemical shift perturbation, *K_d_* = binding dissociation constant, *p* = [protein], and *l* = [ligand].

### Intrinsic tryptophan fluorescence spectroscopy to determine pep13 and pep213 affinity for Fis1

Tryptophan fluorescence data was collected on a PTI Model #814 fluorimeter using a λ_ex_ of 295 nm and λ_em_ of 300 – 400 nm within Starna Cells 3-Q-10 quartz fluorometer rectangular cell with a 10 mm pathlength and excitation/emission slit widths of 4/6 nm, respectively. A concentrated stock of peptides was resuspended in final Fis1 dialysate buffer and a concentration series with the following points was prepared: 0, 1, 3, 7, 10, 30, 70, 100, 300, 700, and 1000 µM peptide. For each titration point, tryptophan emission spectra were collected on samples excluding Fis1 and then, 5 µL of 400 µM Fis1 was added to the sample for a final concentration of 10 µM Fis1 and tryptophan emission spectra were recollected. To account for buffer and tyrosine fluorescence background from the peptide, difference emission spectra were generated by subtracting the background fluorescence intensities from spectra lacking Fis1. The average emission wavelength at each peptide concentration was calculated according to Equation 2 and plotted as a function of the natural log of peptide concentration, which was fit to a Boltzmann sigmoidal model (Equation 3) (Royer *et al*, 1993).

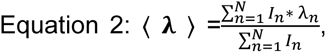

where 〈λ〉 = average emission wavelength, *I_n_* = fluorescence intensity emitted at wavelength λ_n_, and the summation calculated from λ_n_ of 310 to 370 nm.

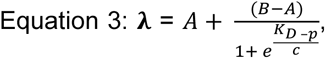

where λ= average emission wavelength, *A* = pre–transition phase, *B* = post–transition phase, *K_D_* = equilibrium dissociation constant, *p* = natural log of the peptide concentration, and *c* = slope of the transition phase. Data were imported into RStudio (Team, 2016) and plotted using Tidyverse (Wickham *et al*, 2019) and readxl (Wickham, 2017)

### Crystallization, data collection, and structure determination of a Fis1-pep213 co-complex

Crystals were grown by using hanging drop vapor diffusion at 19°C. Hanging drops contained a 1.5:1 ratio of a protein-peptide mix (675 µM Fis1ΔTM and 3.375 mM pep213 in 100 mM Hepes, pH 7.4, 200 mM NaCl, 1 mM DTT, 0.02% NaAz) and well-solution (30% PEG 3350) in a 1.25:0.75 ratio with Silver Bullet D5 (Hampton Research HR2-996-41). Droplets were streaked with a 1:1000 dilution of seed crystals (created using Hampton Research ceramic seed bead kit HR4-781) using a cat whisker. Crystals were flash-frozen after rapid-transfer into a cryo-protectant solution (35% PEG 3350) followed by placement in liquid nitrogen. Multiple data sets were collected remotely at the Advanced Photon Source, Argonne National Laboratory (Argonne, IL) LS-CAT beamline 21-ID-F with a MD2-S Microdiffractometer and Rayonix MX300 detector. A total of 180° of data were collected in 0.5° increments with a detector distance of 260 mm. All data were processed using XDS. Molecular replacement using a truncated 1NZN structure was performed with Phenix Phaser-MR followed by an auto-build step using Phenix AutoBuild (Liebschner *et al*, 2019). The crystal belongs to the P4_1_2_1_2 space group with one molecule in the asymmetric unit. Refinement was performed using Phenix.Refine (Liebschner *et al*, 2019) and WinCoot (Emsley *et al*, 2010). Final structure resolution was 1.85 angstroms. Representative images were generated in PyMOL (Schrödinger).

### Treating arterioles with pep213 and measuring endothelial Cell NO bioavailability and endothelium-dependent vasodilation

Vessels from a subset of subjects with and without T2DM were selected at random for these studies. Vessels from healthy subjects were pre-treated with Krebs buffer containing high glucose (33 mM) for six hours and subsequently exposed to 1 µM of TATpep213 or TATscrambled-pep213. Vessels from T2DM subjects were incubated under normal glucose conditions (5 mM) and exposed to either 1 µM of TATpep213 or TATscrambled-pep213. Endothelium-dependent vasodilation with increasing doses of acetylcholine, smooth muscle reactivity to papaverine, and eNOS-dependence of the vasodilatory response to acetylcholine with and without L-NAME were measured as described above and in our previous work (Kizhakekuttu *et al*, 2012; Wang *et al*, 2012; Tanner *et al*, 2017; Widlansky *et al*, 2018; Suboc *et al*, 2013).

### Statistical analyses

Statistical analyses were performed with either GraphPad Prism V7.03 (GraphPad Prism version 8.0.0 for Windows, GraphPad Software, San Diego, California, USA), SigmaPlot Version 12.5 (SysTATSoftware, San Jose, California, USA), or RStudio (version 4.0.4). P values <0.05 were considered statistically significant. Data were represented as box and whisker plots when appropriate, which present data as the median, a 25^th^-75^th^ percentile box, and whiskers extending to 1.5 times the inter-quartile range. All other data are presented as mean ± SEM unless otherwise stated. All functional vessel data was analyzed by two-way ANOVA with post-hoc testing to determine the differences between groups (Dunnett’s multiple comparison test) and dose-response (Tukey’s multiple comparison tests). The TEER assay data and bioenergetics data were also analyzed by two-way ANOVA followed by post-hoc testing (Tukey’s multiple comparison tests) to determine the differences between normal, low, and high glucose treatments. DAF-2 DA fluorescence intensity in human vessels, Fis1 knockdown-efficiency in HMEC-1 cells, and measurements of mitochondrial protein amounts by Western blot were analyzed by one-way ANOVA followed by Tukey’s multiple comparisons to assess differences between groups. DAF-2 DA fluorescence intensity in HMEC-1 cells transfected with siFis1 and scrambled siRNA were analyzed by two-way ANOVA followed by post-hoc testing (Tukey’s multiple comparison tests). NO production and relative ratio of P-eNOS and β-actin in HMEC-1 cells stimulated with A23187 were analyzed using paired Student’s t-test.

## Data Availability

All R scripts used for data analysis and visualization are available upon request and/or for download at https://github.com/Hill-Lab/. Raw data is available upon request.

## Acknowledgement

We gratefully acknowledge Ms. Cathy Paddock and Dr. Peter Newman for training and access to the ECIS instrument for TEER measurements. We also acknowledge KAN’s cat, Winnie, for her whisker used in seeding the crystallography trays. This research used resources of the Advanced Photon Source; a U.S. Department of Energy (DOE) Office of Science User Facility operated for the DOE Office of Science by Argonne National Laboratory under Contract No. DE-AC02-06CH11357. Use of the LS-CAT Sector 21 was supported by the Michigan Economic Development Corporation and the Michigan Technology Tri-Corridor (Grant 085P1000817).

## Author Contributions

KAN: Conceptualization, methodology, software, validation, formal analysis, investigation, writing (original, review, editing), visualization

MK: Methodology, investigation, analyses, writing (original, review, editing)

JME: Conceptualization, methodology, software, validation, formal analysis, investigation, writing (original), visualization

JW: Methodology, investigation, analyses, writing (reviewing, editing)

MCH: Conceptualization, formal analysis, visualization writing (review, editing) VKP: Methodology, investigation, writing (reviewing, editing)

BCH: Methodology, investigation, writing (reviewing, editing) LAA: Supervision, writing (reviewing, editing)

DZT: Investigation, formal analysis. FCP: Validation, formal analysis MLR: Investigation, formal analysis. DMJ: Investigation, formal analysis.

RBH: conceptualization, supervision, validation, funding acquisition, writing (review, editing)

MEW: conceptualization, supervision, funding acquisition, validation, resources, writing (review, editing)

## The Paper Explained

### Problem

Vascular endothelial dysfunction is a significant precursor for the development of microvascular and macrovascular diseases including those found in diabetic patients. Hyperglycemia stimulates mitochondrial dysfunction with increased reactive oxygen species (ROS) generation and excess mitochondrial network fragmentation, all of which drive endothelial dysfunction. Attempts to utilize antioxidant-based therapeutics to negate the increased ROS have failed in clinical trials, suggesting an alternative approach is necessary. Whether targeting excess mitochondrial network fragmentation can improve endothelial dysfunction in diabetic patients remains unknown.

### Results

Using an ex vivo system, we determined that mitochondrial fission 1 protein (Fis1) appears to play a causative role in diabetic endothelial dysfunction as its overexpression in healthy vessels decreases their ability to vasodilate. Conversely, genetic silencing of Fis1 in diabetic vessels, or healthy vessels exposed to abnormal glucose concentrations, significantly improved endothelium-dependent vasodilation of arterioles and decreased excess mitochondrial ROS generation. Given these results, we developed a first-in-class Fis1 inhibitor which induced similar beneficial effects in human arterioles.

### Impact

Our results suggest that Fis1 plays a critical role in the development of diabetic endothelial dysfunction. These effects appear to be mediated through detrimental increases in mitochondrial ROS and can be rescued with a pharmacological Fis1 inhibitor. Ultimately, this work identifies a novel mechanism for treating diabetic microvascular disease via pharmacological inhibition of Fis1.

