## Supplemental Data and Tables for "A novel Fis1 inhibiting peptide reverses diabetic endothelial dysfunction in human resistance arteries"

**Supplementary Materials**

*Supplemental Figures*

**Supplemental Figure 1.** Genetic manipulation of Fis1 levels significantly alters total Fis1 without altering vessel response to papaverine. **A.** Maximum vasodilation of human vessels due to papaverine stimulation scrambled and Fis1 siRNA treatment in diabetic vessels (DM), vessels overexpressing Fis1 due to lentiviral transfection (Fis1 OE), and siRNA transfected healthy vessels exposed to high (HG, 33 mM) or low (LG, 2.5 mM) glucose levels reveal negligible changes in smooth muscle relaxation secondary to siRNA treatment or Fis1 over-expression (all comparisons not statistically significant except for scrambled and Fis1 siRNA treatment in DM vessels (94.1±3,3% vs. 98.6±1.6% for scrambled control siRNA and Fis1 siRNA, respectively, n=6, p=0.04) **B.** Lentiviral transfection of Fis1 in healthy small resistance vessels induces a significant increase in total Fis1 protein content as determined by fluorescence intensity normalized to total sample protein content using Wes (n=4, p=0.0008 by Student’s t-test). **C.** Fis1 siRNA treatment in small resistance vessels from diabetic patients significantly decreases total Fis1 protein levels as determined by Wes (n=5, p=0.0044 by Student’s t-test). The boxes represent 25^th^ and 75^th^ percentiles, with the horizontal line representing the median fluorescence intensity.

**Supplemental Figure 2.** Genetic silencing of Fis1 increases NO levels in healthy vessels exposed to low (2.5 mM) and high (33 mM) glucose conditions. Human resistance vessels isolated from normal healthy subjects were treated with Fis1 siRNA and the fluorescent NO marker DAF-2 DA was measured under low (**A**, 2.5 mM, 2 hrs) or high (**B**, 33 mM, 6 hrs) glucose conditions upon Fis1 siRNA treatment. For low glucose conditions, **A**, n=8, overall p=0.01; p=0.03 for control siRNA vs. Fis1 siRNA; p=0.003 for control siRNA+L-NAME vs Fis1 siRNA; p=0.03 for Fis1 siRNA vs Fis1 siRNA+L-NAME. For high glucose conditions, **B**, n=9, overall p=0.04; p=0.01 for scrambled siRNA vs Fis1 siRNA; p=0.03 for Fis1 siRNA vs control siRNA+L-NAME and p=0.047 for Fis1 siRNA vs Fis1 siRNA+L-NAME. The boxes represent 25th and 75th percentiles, with the horizontal line representing the median fluorescence intensity. Statistical significance was determined by ANOVA with post-hoc Tukey’s.

**Supplemental Figure 3.** Genetic silencing of Drp1 protects against glucose-induced impairments in vasodilation while concurrently improving NO bioavailability. **A.** Genetic silencing of Drp1 via Drp1 siRNA treatment protected against reductions in nitric oxide (NO) bioavailability in healthy human arterioles exposed to high glucose (33mM): n=9, p=0.02 overall; p<0.05 for Drp1 siRNA vs. all other treatments and **B.** low glucose (2.5mM): n=5; overall p=0.003, p<0.03 for Drp1 siRNA vs. all other treatments. The boxes represent 25^th^ and 75^th^ percentiles, with the horizontal line representing the median fluorescence intensity. **C.** Transfection with Drp1 siRNA protected against high glucose-induced impairment in endothelium-dependent vasodilation (n=6, p<0.0001 overall, p≤0.0001 at indicated Ach doses for control siRNA vs. Drp1 siRNA). L-NAME abolished this improvement of vasodilation under Drp1 knockdown conditions (p<0.0001 for Drp1 siRNA vs. Drp1 siRNA + L-NAME). **D.** Suppression of Drp1 expression in arterioles from humans with T2DM using Drp1 siRNA trended towards reverse impaired endothelium-dependent vasodilation (n=4, P=0.076). Data are mean ± SEM. Statistical significance determined by ANOVA with post-hoc Tukey’s.

**Supplemental Figure 4.** Suppressing Fis1 in endothelial cells improves NO bioavailability and barrier function without altering cellular energetics. **A.** Fis1 knockdown efficiency in HMEC-1 cells with siRNA treatment. Fis1 levels were significantly reduced (n=3, overall p<0.0001) in HMEC-1 cells transfected with Fis1 siRNA Fis1 compared to cells transfected with scrambled control siRNA (p=0.0002) and non-transfected HMEC-1 cells (p=0.0001). **B.** Suppression of Fis1 expression under low (2.5mM) and high (33 mM) glucose conditions improves steady-state junction stability in endothelial cell monolayers. Comparison of electric cell-substrate impedance sensing measurements (ECIS) in human microvascular endothelial cell (HMEC-1) transfected with Fis1 siRNA and control siRNA between normal (5 mM), low (2.5 mM), and high (33 mM) glucose conditions. (n=4 for each treatment, p<0.001 overall for control siRNA vs. Fis1 siRNA for both high and low glucose studies. *p<0.05, **p<0.01, ***p<0.001, ****p<0.0001. Measures of extracellular acidification rate (ECAR, n=5) (**C**) and oxygen consumption rate (OCR, n=5) (**D**) in human microvascular endothelial cells (HMEC-1) transfected with Fis1 siRNA and pre-incubated with normal (5mM for 2 hours), low (2.5 mM for 2 hours) or high (33 mM for 6 hours) glucose conditions. Wild-type (WT) cells represent HMEC-1 cells not transfected and grown under normal glucose conditions. ECAR and OCR were measured under basal conditions followed by the sequential addition of the indicated compounds. No statistically significant differences were present between the treatments. Data is mean ± SD. **E.** HMEC-1 cells were treated with vehicle control (basal condition) or 5 μM A23187 for 2 hours, followed by a 15-minute incubation with DAF-2 DA (5µM) prior to measurement of DAF-2 DA fluorescence intensity (n=7, p<0.0001 overall; basal Fis1 siRNA condition vs. stimulated Fis1 siRNA p<0.0001; basal control siRNA condition vs. stimulated Fis1 siRNA condition p=0.0003; stimulated Fis1 siRNA condition vs. stimulated control siRNA p=0.02). **F.** Representative western blot probing for p-eNOS(Ser1177) and β-actin with and without the addition of 5 μM A23187. **G.** A quantitative measure of p-eNOS-(Ser1177) treated with and without 5 μM A23187 shows a significant increase in p-eNOS(Ser1177) upon ionophore stimulation (n=6, p=0.03 by Student’s t-test). **H.** NO production is significantly increased as measured using DAF-2 DA (5 μM) upon the addition of A23187 (n=4, p=0.02 by Student’s t-test). Boxes represent 25^th^ and 75^th^ percentiles, with the horizontal line representing the median. Statistical significance determined by ANOVA with post-hoc Tukey’s unless otherwise indicated.

**Supplemental Figure 5.** Molecular inhibition of Fis1 under high (33mM) and low (2.5mM) glucose conditions did not alter the expression of other mitochondrial proteins. The expression of selected mitochondrial proteins was measured from immortalized HMEC-1 cells transfected with siRNA Fis1 or scrambled siRNA and pre-incubated with different glucose conditions: high glucose (HG, 33mM for six hours), normal glucose (NG, 5mM for 2 hours) and low glucose (LG, 2.5mM for 2 hours). The expression of each protein was normalized to total protein across the samples. (n=4-10 for individual proteins). While there are differences in the expression of some mitochondrial proteins seen when comparing different glucose concentration exposures, knockdown of Fis1 expression with siRNA did not affect the expression of any mitochondrial protein except for Fis1. Statistically significant differences determined by ANOVA with post-hoc Tukey’s and indicated by * for p<0.05, ** for p<0.01, *** for p<0.001, and **** for p<0.0001.

**Supplemental Figure 6.** Fis1 binds to pep2 and pep13, but not a TAT-fusion randomized sequence of pep213. ^1^H, ^15^N HSQC spectral overlays of 50 µM ^15^N-Fis1^1-125^ with increasing amounts (0-2mM) unlabeled pep2 (**A**) or pep13 (**B**) with changes in chemical shift indicative of binding. **C.** Affinity determination of pep13 for Fis1 using intrinsic tryptophan fluorescence in which increasing pep13 (0-1mM) was titrated into 10 µM Fis1 and the resulting average emission wavelength was fit to determine an apparent *K_D_* = 12.1 ± 4.1 µM. Residuals to the fits shown on the right. **D.**  ^1^H, ^15^N HSQC spectral overlays of 50 µM ^15^N-Fis1^1-125^ with increasing amounts (0-2mM) an unlabeled TATpep213 construct with a randomized (scrambled) peptide sequence with minimal changes in chemical shift at 2 mM peptide, indicative of very weak to no binding. **E.** Zoomed in region from spectra in (**D**).

**Supplemental Figure 7.** Pep213 modeling with surrounding electron density after molecular replacement and pre- and post-refinement. Pre-refinement map shown at 2σ and post-refinement map shown at 1σ.

*Supplemental Methods*

**siRNA sequences**

5’ to 3’ sequences for siRNA to Fis1 and Drp1 were the following:

Fis1:

A: rGrGrUrGrCrGrGrArGrCrArArGrUrArCrArArUrGrArUrGAC

B: rArCrUrArCrCrGrGrCrUrCrArArGrGrArArUrArCrGrArGAA

C: rArCrArGrUrArGrArCrUrGrUrArGrUrGrUrGrArGrGrCrUCG

Drp1:

A: rArGrArGrUrGrUrArArCrUrGrArUrUrCrArArUrCrCrGrUGA

B: rArGrGrArUrArUrUrGrArGrCrUrUrCrArArArUrCrArGrAGA

C: rCrCrCrUrUrArArArCrUrGrArGrUrCrArArGrArUrCrUrGAA

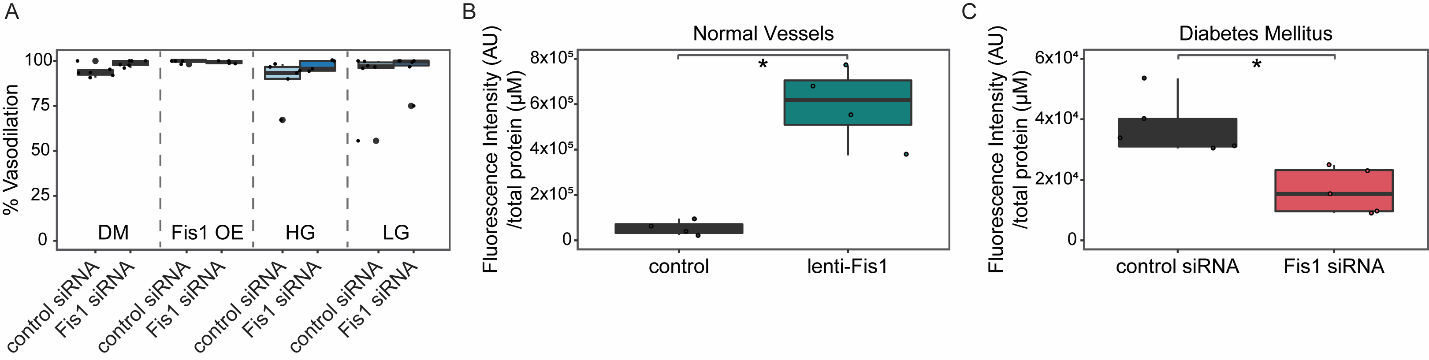

**Supplemental Figure 1.** Genetic manipulation of Fis1 levels significantly alters total Fis1 without altering vessel response to papaverine.

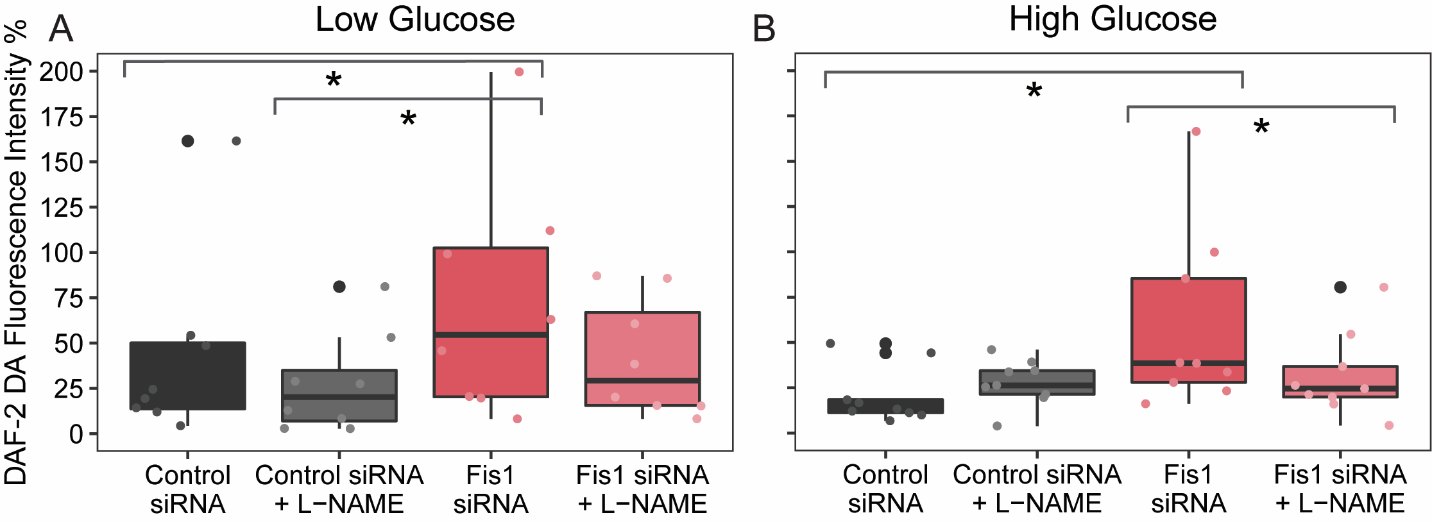

**Supplemental Figure 2.** Genetic silencing of Fis1 increases NO levels in healthy vessels exposed to low (2.5 mM) and high (33 mM) glucose conditions.

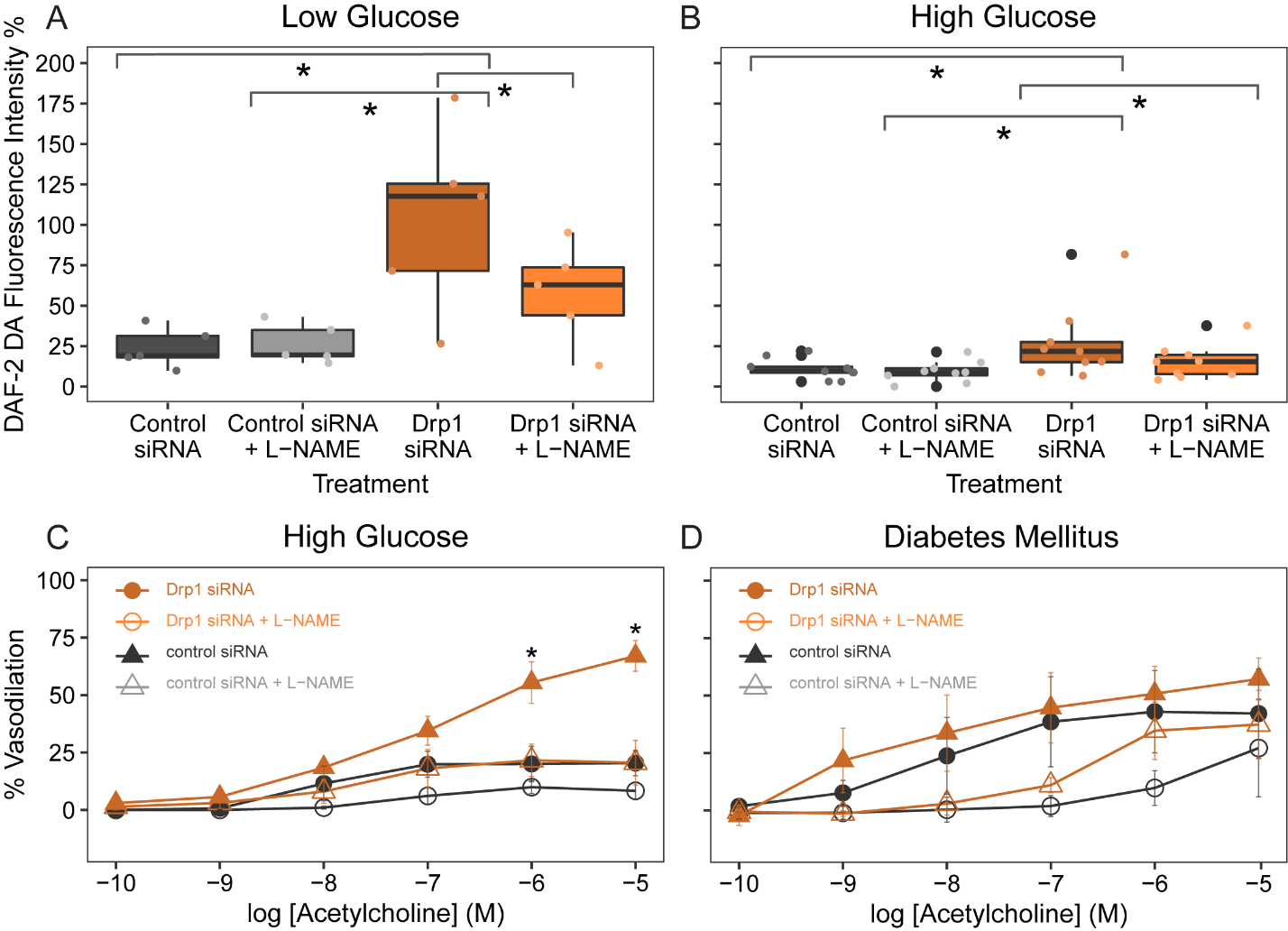

**Supplemental Figure 3.** Genetic silencing of Drp1 protects against glucose-induced impairments in vasodilation while concurrently improving NO bioavailability.

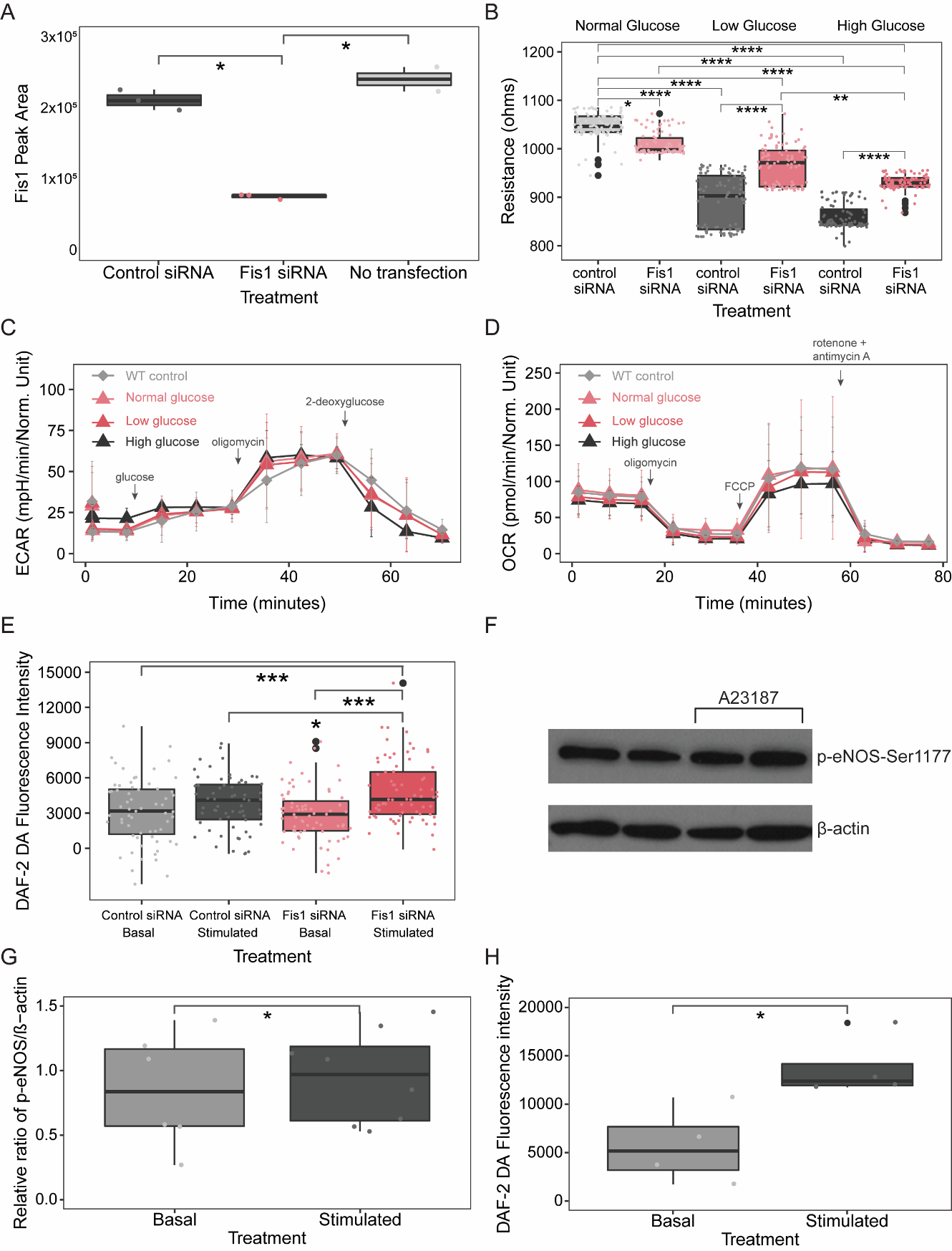

**Supplemental Figure 4.** Suppressing Fis1 in endothelial cells improves NO bioavailability and barrier function without altering cellular energetics.

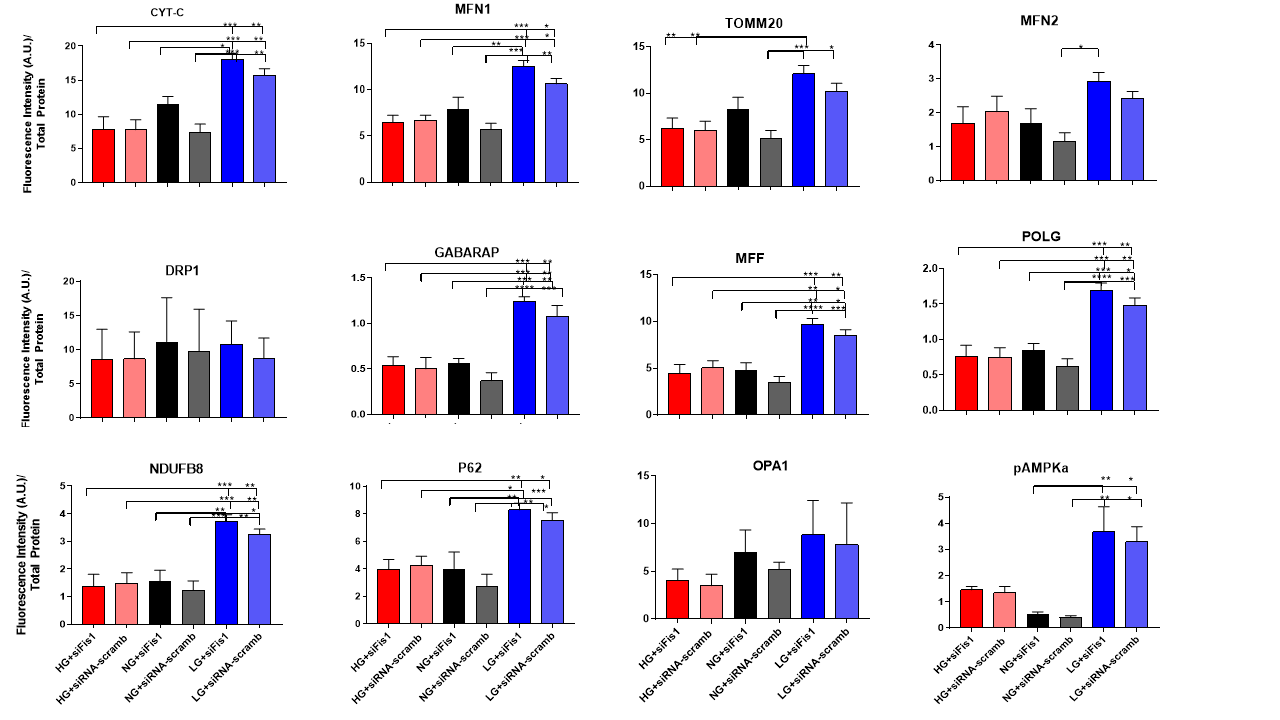

**Supplemental Figure 5.** Molecular inhibition of Fis1 under high (33mM) and low (2.5mM) glucose conditions did not alter the expression of other mitochondrial proteins.

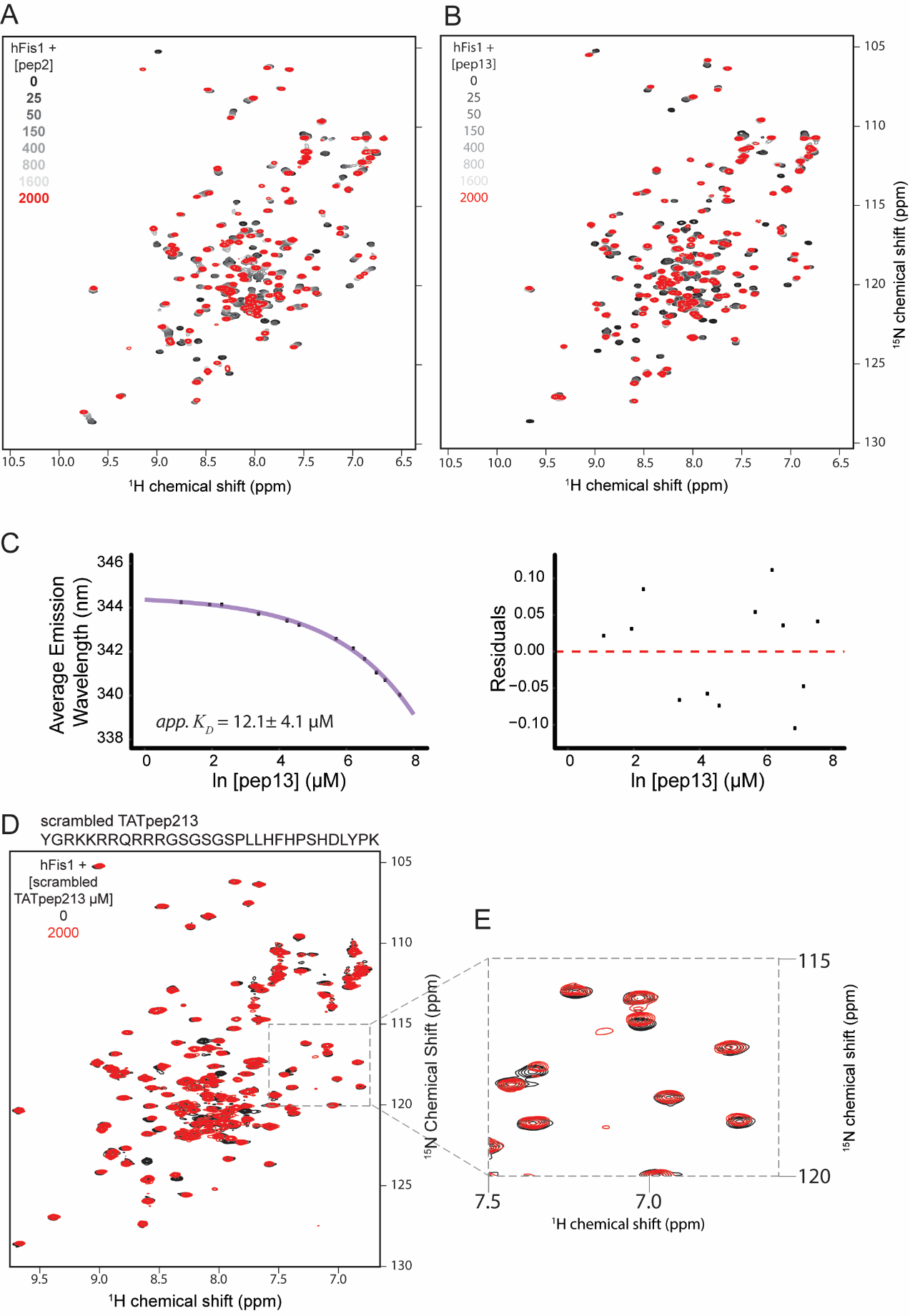

**Supplemental Figure 6.** Fis1 binds to pep2 and pep13, but not a TAT-fusion randomized sequence of pep213.

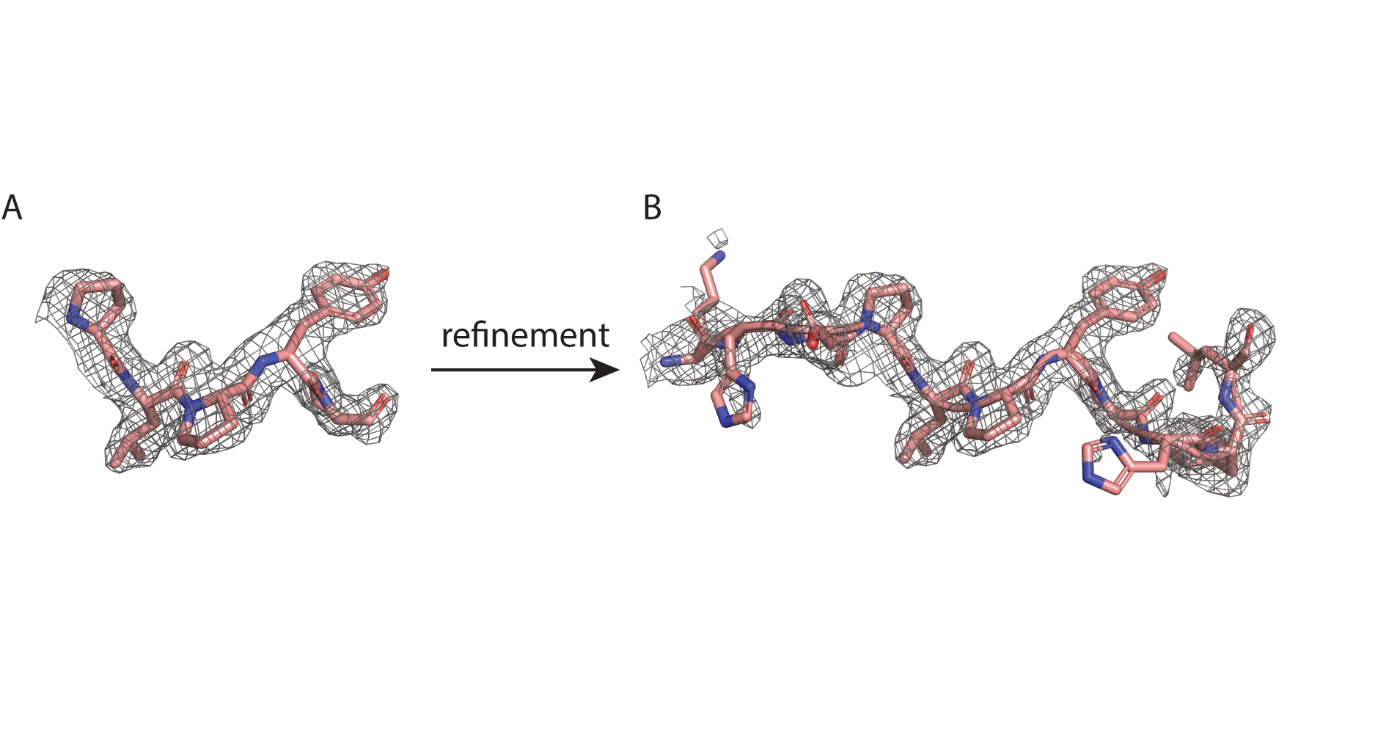

**Supplemental Figure 7.** Pep213 modeling with surrounding electron density after molecular replacement and pre- and post-refinement.

**Supplemental Tables**

| Structure | Fis1-Pep213 Co-complex |
| --- | --- |
| Resolution range | 37.57 - 1.851 (1.917 - 1.851) |
| Space group | P 41 21 2 |
| Unit cell | 54.798 54.798 103.198 90 90 90 |
| *Data Processing* |  |
| Total reflections | 192076 (16540) |
| Unique reflections | 14024 (1342) |
| Redundancy | 13.7 (12.3) |
| Completeness (%) | 99.72 (98.53) |
| Mean I/σ(I) | 23.77 (1.53) |
| R_merge_ (%) | 6.40 |
| R_meas_ (%) | 6.66 |
| R_pim_ (%) | 1.80 |
| *Refinement* |  |
| Reflections used in refinement | 14023 (1343) |
| R_work_ (%) | 20.81 (28.18) |
| R_free_ (%) | 25.92 (38.09) |
| Number of non-hydrogen atoms | 1201 |
| Macromolecules/solvent | 1116/85 |
| RMS(bonds)/(angles) | 0.006/0.79 |
| Average B-factor/macromolecules/solvent | 40.82/40.61/43.62 |
| *Ramachandran, rotamer, and clash statistics* |  |
| Ramachandran favored (%) | 96.97 |
| Ramachandran allowed (%) | 3.03 |
| Ramachandran outliers (%) | 0.00 |
| Rotamer outliers (%) | 0.00 |
| Clashscore | 3.11 |

Statistics for the highest-resolution shell are shown in parentheses.

**Supplemental Table 1.** Fis1-pep213 co-complex structure data collection and refinement statistics.

|  | Non-Diabetic  (LG; n=6) | Non-Diabetic  (HG; n=5) | Type 2 DM (n=6) | p-value |
| --- | --- | --- | --- | --- |
| Age | 35±17 | 58±5 | 61±10 | 0.01 |
| Sex (#female) | 3 | 3 | 0 | 0.12 |
| Smoking Status (#never) | 1 | 1 | 3 | 0.03 |
| History of Hypertension | 0 | 0 | 5 | 0.0002 |
| History of High Cholesterol | 0 | 0 | 5 |  |
| Body mass Index (kg/m2) | 28±5 | 26±3 | 29±5 | 0.80 |
| Waist Circumference (cm) | 95±18 | 88±10 | 109±4 | 0.18 |
| Systolic Blood Pressure (mmHg) | 120±17 | 126±11 | 126±11 | 0.69 |
| Diastolic Blood Pressure (mmHg) | 72±10 | 74±11 | 74±8 | 0.85 |
| Fasting Glucose (mg/dL) | 83±11 | 82±7 | 174±35 | <0.0001 |
| Hemoglobin A1C (%) | 5.3±0.3 | 5.4±0.2 | 8.3±1.8 | 0.0009 |
| Creatinine (mg/dL) | 0.82±0.16 | 0.92±0.18 | 0.95±0.19 | 0.25 |
| Total Cholesterol (mg/dL) | 204±33 | 195±16 | 174±41 | 0.35 |
| HDL Cholesterol | 71±15 | 86±19 | 49±6 | 0.009 |
| LDL Cholesterol (mg/dL) | 115±34 | 95±18 | 84±31 | 0.21 |
| Medications (% on Therapy) |  |  |  |  |
| Biguanide | 0 | 0 | 83 |  |
| Sulfonylurea | 0 | 0 | 83 |  |
| Thiazoladinedione | 0 | 0 | 0 |  |
| DPP4 inhibitor | 0 | 0 | 33 |  |
| GLP-1 Agonist | 0 | 0 | 0 |  |
| Insulin | 0 | 0 | 0 |  |
| HMG CoA reductase Inhibitor | 0 | 0 | 100 |  |
| ACE Inhibitor | 0 | 0 | 83 |  |
| Angiotensin II Receptor Blocker | 0 | 0 | 17 |  |
| GLP1 Agonist | 0 | 0 | 17 |  |
| SGLT2 Inhibitor | 0 | 0 | 0 |  |

**Supplemental table 2**. Demographic, clinical, and medication information and *in vivo* vascular function for 16 subjects with resistance arterioles that underwent Fis1 siRNA transfection and studied for vasodilation in response to acetylcholine.

|  | Non-DM  (LG; n=8) | Non-DM  (HG; n=9) | p-value |
| --- | --- | --- | --- |
| Age | 42±16 | 35±14 | 0.54 |
| Sex (#female) | 5 | 8 | 0.35 |
| Smoking Status (#never) | 3 | 2 | 0.38 |
| History of Hypertension | 0 | 0 | - |
| History of High Cholesterol | 0 | 0 | - |
| Body mass Index (kg/m2) | 23±3 | 23±6 | 0.89 |
| Waist Circumference (cm) | 85±11 | 82±17 | 0.78 |
| Systolic Blood Pressure (mmHg) | 125±9 | 111±12 | 0.02 |
| Diastolic Blood Pressure (mmHg) | 68±5 | 68±12 | 0.82 |
| Fasting Glucose (mg/dL) | 85±3 | 89±9 | 0.50 |
| Hemoglobin A1C (%) | 5±0.2 | 5.2±0.3 | 0.27 |
| Creatinine (mg/dL) | 0.8±0.1 | 0.74±0.11 | 0.12 |
| Total Cholesterol (mg/dL) | 197±31 | 171±22 | 0.14 |
| HDL Cholesterol | 70±16 | 63±23 | 0.81 |
| LDL Cholesterol (mg/dL) | 109±23 | 89±22 | 0.12 |

**Supplemental table 3.** Demographic, clinical, and medication information and in vivo vascular function for 17 subjects with resistance arterioles that underwent Fis1 siRNA transfection and studied for biological NO production using DAF2DA fluorescence.

|  | Non-Diabetic (HG; n=5) | Type 2 DM (n=4) | p-value |
| --- | --- | --- | --- |
| Age | 33±11 | 46±15 | 0.43 |
| Sex (#female) | 2 | 3 | 0.29 |
| Smoking Status (#never) | 5 | 2 | 0.49 |
| History of Hypertension | 0 | 3 | 0.01 |
| History of High Cholesterol | 0 | 2 | 0.20 |
| Body mass Index (kg/m2) | 27±7 | 30±5 | 0.49 |
| Waist Circumference (cm) | 89±19 | 95±12 | 0.75 |
| Systolic Blood Pressure (mmHg) | 119±20 | 131±33 | 0.96 |
| Diastolic Blood Pressure (mmHg) | 74±15 | 79±22 | 0.95 |
| Fasting Glucose (mg/dL) | 71±9 | 142±55 | 0.002 |
| Hemoglobin A1C (%) | 5.2±0.5 | 8.4±1.0 | <0.0001 |
| Creatinine (mg/dL) | 0.89±0.23 | 0.82±0.22 | 0.40 |
| Total Cholesterol (mg/dL) | 179±18 | 163±16 | 0.09 |
| HDL Cholesterol | 74±27 | 56±3 | 0.39 |
| LDL Cholesterol (mg/dL) | 87±13 | 90±19 | 0.40 |
| Medications (% on Therapy) |  |  |  |
| Biguanide | 0 | 75 |  |
| Sulfonylurea | 0 | 25 |  |
| Thiazoladinedione | 0 | 0 |  |
| DPP4 inhibitor | 0 | 0 |  |
| GLP-1 Agonist | 0 | 0 |  |
| Insulin | 0 | 25 |  |
| HMG CoA reductase Inhibitor | 0 | 50 |  |
| ACE Inhibitor | 0 | 25 |  |
| Angiotensin II Receptor Blocker | 0 | 0 |  |
| SGLT2 Inhibitor | 0 | 0 |  |

**Supplemental table 4.** Demographic, clinical, and medication information and in vivo vascular function for 13 subjects with resistance arterioles that underwent Drp1 siRNA transfection and studied for vasodilation in response to acetylcholine.

|  | Non-Diabetic (LG; n=5) | Type 2 DM  (HG; n=9) | p-value |
| --- | --- | --- | --- |
| Age | 29±9 | 42±14 | 0.27 |
| Sex (#female) | 3 | 6 | 0.37 |
| Smoking Status (#never) | 3 | 2 | 0.36 |
| History of Hypertension | 0 | 0 | na |
| History of High Cholesterol | 0 | 0 | na |
| Body mass Index (kg/m2) | 25±6 | 29±6 | 0.26 |
| Waist Circumference (cm) | 86±16 | 94±15 | 0.70 |
| Systolic Blood Pressure (mmHg) | 116±10 | 126±12 | 0.61 |
| Diastolic Blood Pressure (mmHg) | 71±11 | 71±10 | 0.91 |
| Fasting Glucose (mg/dL) | 82±13 | 87±9 | 0.45 |
| Hemoglobin A1C (%) | 5.1±0.1 | 5.2±0.3 | 0.42 |
| Creatinine (mg/dL) | 0.81±0.15 | 0.86±0.21 | 0.53 |
| Total Cholesterol (mg/dL) | 170±35 | 178±27 | 0.72 |
| HDL Cholesterol | 67±10 | 61±12 | 0.74 |
| LDL Cholesterol (mg/dL) | 86±29 | 97±20 | 0.76 |

**Supplemental table 5.** Demographic, clinical, and medication information and in vivo vascular function for 16 subjects with resistance arterioles that underwent Drp1 siRNA transfection and studied for biological NO production using DAF2DA fluorescence.

|  | Non-DM  (HG; n=6) | T2DM (n=4) | p-value |
| --- | --- | --- | --- |
| Age | 35±14 | 46±15 | 0.02 |
| Sex (#female) | 1 | 2 | 0.29 |
| Smoking Status (#never) | 5 | 1 | 0.49 |
| History of Hypertension | 0 | 3 | 0.01 |
| History of High Cholesterol | 0 | 3 | 0.20 |
| Body mass Index (kg/m2) | 27±7 | 30±5 | 0.007 |
| Waist Circumference (cm) | 89±19 | 95±12 | 0.04 |
| Systolic Blood Pressure (mmHg) | 117±7 | 125±16 | 0.32 |
| Diastolic Blood Pressure (mmHg) | 70±5 | 77±7 | 0.10 |
| Fasting Glucose (mg/dL) | 92±6 | 129±36 | 0.04 |
| Hemoglobin A1C (%) | 5.0±0.3 | 8.0±2.4 | 0.01 |
| Creatinine (mg/dL) | 0.92±0.16 | 0.77±0.07 | 0.11 |
| Total Cholesterol (mg/dL) | 179±35 | 179±55 | 0.98 |
| HDL Cholesterol | 55±19 | 50±18 | 0.67 |
| LDL Cholesterol (mg/dL) | 110±22 | 104±41 | 0.79 |
| Medications (% on Therapy) |  |  |  |
| Biguanide | 0 | 100 |  |
| Sulfonylurea | 0 | 50 |  |
| Thiazoladinedione | 0 | 0 |  |
| GLP-1 Agonist | 0 | 25 |  |
| DPP4 inhibitor | 0 | 0 |  |
| Insulin | 0 | 25 |  |
| HMG CoA reductase Inhibitor | 0 | 50 |  |
| ACE Inhibitor | 0 | 25 |  |
| Angiotensin II Receptor Blocker | 0 | 25 |  |
| SGLT2 Inhibitor | 0 | 0 |  |

**Supplemental table 6.** Demographic, clinical, and medication information and in vivo vascular function for 10 subjects with resistance arterioles exposed to pep213 and studied for vasodilation in response to acetylcholine.

|  | Non-DM  (HG; n=5) | T2DM (n=4) | p-value |
| --- | --- | --- | --- |
| Age | 54±19 | 46±15 | 0.41 |
| Sex (#female) | 4 | 2 | 0.29 |
| Smoking Status (#smoker) | 0 | 0 | - |
| History of Hypertension | 1 | 1 | 0.72 |
| History of High Cholesterol | 0 | 1 | 0.56 |
| Body mass Index (kg/m2) | 29±6 | 35±10 | 0.27 |
| Medications (% on Therapy)  ACE Inhibitor | 0 | 0 | - |
| Angiotensin II Receptor Blocker | 0 | 1 | 0.56 |

**Supplemental table 7.** Demographic, clinical, and medication information and in vivo vascular function for 9 subjects with resistance arterioles exposed to TATpep213and scrambled peptide-TAT and studied for vasodilation in response to acetylcholine.

|  | Fis1 Over-Expression vs. Control (N=4) | Fis1 Over-Expression with and without pep213 exposure vasoactivity and Fis1 Over-Expression MitoNeoD Studies (n=5) |
| --- | --- | --- |
| Age | 42±5 | 41±4 |
| Sex (#female) | 3 | 5 |
| Smoking Status (#smoker) | 0 | 0 |
| History of Hypertension | 0 | 0 |
| History of High Cholesterol | 0 | 0 |
| Body mass Index (kg/m2) | 34±4 | 27±3 |

**Supplemental table 8.** Demographic and clinical information and in vivo vascular function and MitoNeoD studies the following experiments: (1) Fis1 over-expression vs. control transfection vasoactivity studies (N=4), (2) Fis 1 over-expression with exposure to pep213-tat vs. scrambled peptide-tat and studied for vasodilation in response to acetylcholine and MitoNeoD fluorescence intensity.
